# The mitochondrial carrier SFXN1 is critical for Complex III integrity and cellular metabolism

**DOI:** 10.1101/2020.06.18.157495

**Authors:** Michelle Grace Acoba, Ebru S. Selen Alpergin, Santosh Renuse, Lucía Fernández-del-Río, Ya-Wen Lu, Catherine F. Clarke, Akhilesh Pandey, Michael J. Wolfgang, Steven M. Claypool

## Abstract

Mitochondrial carriers (MC) mediate the passage of small molecules across the inner mitochondrial membrane (IMM) enabling regulated crosstalk between compartmentalized reactions. Despite MCs representing the largest family of solute carriers in mammals, most have not been subjected to a comprehensive investigation, limiting our understanding of their metabolic contributions. Here, we functionally characterized SFXN1, a member of the non-canonical, sideroflexin MC family. We find that SFXN1, an integral membrane protein in the IMM with an uneven number of transmembrane domains, is a novel TIM22 substrate. SFXN1 deficiency specifically impairs Complex III (CIII) biogenesis, activity, and assembly, compromising coenzyme Q levels. This CIII dysfunction is independent of one-carbon metabolism, the known primary role for SFXN1 as a mitochondrial serine transporter. Instead, SFXN1 supports CIII function by participating in heme and central carbon metabolism. Our findings highlight the multiple ways that SFXN1-based amino acid transport impacts mitochondrial and cellular metabolic efficiency.

## INTRODUCTION

Mitochondria are classically viewed as energy-producing organelles. Lining the folds of the inner mitochondrial membrane (IMM) cristae is the respiratory chain —a group of multimeric complexes (Complex I-IV) and mobile electron carriers— that facilitates the transfer of electrons from NADH and FADH_2_ to O_2_ while pumping protons to the intermembrane space (IMS). The use of the proton gradient powers the ATP synthase (Complex V), whose rotary action creates ATP by oxidative phosphorylation (OXPHOS) (Abrahams et al., 1994; Hutton and Boyer, 1979; Mitchell, 1961). Because this is the major source of cellular ATP, perturbations in OXPHOS underlie various mitochondrial pathologies including rare syndromes and a growing number of aging-related diseases (Sun et al., 2016; Vafai and Mootha, 2012).

Beyond a role in bioenergetics, mitochondria participate in the biosynthesis of nucleotides, lipids, amino acids, and numerous metabolites. The vital position of mitochondria in cells is underscored by the emerging appreciation that they contribute to intracellular signaling and regulatory networks (Spinelli and Haigis, 2018). At the core of this are the mitochondrial carriers (MC) that coordinate the transport of diverse solutes into and out of mitochondria. Collectively, MCs enable mitochondria to communicate with the rest of the cell to achieve homeostasis and allow this complex organelle to influence metabolic plasticity, an attribute that is paramount for survival in a constantly changing metabolic landscape (Palmieri and Monne, 2016; Taylor, 2017).

Dysfunction in MCs result in pathologies affecting different organ systems and manifesting at varying severities (Ogunbona and Claypool, 2019). Many of the disorders involve OXPHOS defects while others lead to specific metabolic problems. For example, deficiencies in the adenine nucleotide translocase (ANT) isoform 1 and the phosphate carrier (PiC) cause myopathy and hypertrophic cardiomyopathy (Bakker et al., 1993; Bhoj et al., 2015; Echaniz-Laguna et al., 2012; Korver-Keularts et al., 2015; Mayr et al., 2007; Palmieri et al., 2005; Thompson et al., 2016). On the other hand, mutations in *SLC25A38*, which code for a glycine transporter, impinge on heme production and are associated with sideroblastic anemia (Guernsey et al., 2009; Harigae and Furuyama, 2010; Horvathova et al., 2010). Numerous MCs have also been implicated in cancer progression, including the understudied amino acid transporters that support biomass production for increased cell proliferation (Lytovchenko and Kunji, 2017).

To gain further insight into the functional organization of MCs, recent work by our group used a proteomic approach to systematically map the binding partners of human ANT1 and ANT2 (Lu et al., 2017)—the archetypal members of the solute carrier 25 (SLC25) family—that mediate the exchange of ADP and ATP across the IMM (Klingenberg, 2008; Pebay-Peyroula et al., 2003). Among the interactors is another group of mitochondrial membrane proteins—the sideroflexin (SFXN) family, an enigmatic group whose function is only now emerging. SFXN1-5 are categorized under SLC56, a group separate from SLC25, because they are predicted to possess four to five transmembrane domains (TMDs) (Fleming et al., 2001), in contrast to the canonical view that MCs consist of six, forming a structure with three-fold pseudosymmetry (Palmieri, 2004). SFXN1, the founding member of the SLC56 family, was identified by positional cloning of the mutation in the *flexed-tail* mouse model with sideroblastic anemia (Fleming et al., 2001), although its role as the causal mutation has since come into question (Hegde et al., 2007; Lenox et al., 2005). SFXN1 was recently identified as a mitochondrial serine transporter essential for one-carbon (1C) metabolism (Kory et al., 2018), a process wherein folate species are generated and used in biosynthetic pathways required for cell proliferation (Ducker and Rabinowitz, 2017). Notably, while its role as a serine transporter was shown, competition assays indicate that SFXN1 can transport additional amino acids. Mutations in another isoform, SFXN4, are associated with mitochondriopathy and macrocytic anemia (Hildick-Smith et al., 2013). However, the nature and extent of how SFXNs impact mitochondrial biology is still essentially unexplored.

Here, we investigated SFXN1 in mammalian cells. We found that in HEK293, SFXN1 is the most abundant isoform among the SFXNs and demonstrate that it is an integral polytopic protein in the IMM whose topology is consistent with its five predicted TMDs. The steady-state protein abundance of SFXNs is reduced without acylglycerol kinase (AGK), a metazoan-specific component of the TIM22 translocon crucial for the biogenesis of six TMD-containing MCs (Kang et al., 2017; Vukotic et al., 2017); this indicates SFXNs are novel TIM22 substrates. Although predicted to be involved in the mitochondrial transport of a component necessary for iron usage (Fleming et al., 2001), SFXN1 does not govern iron homeostasis in HEK293. Rather, we find that Complex III (CIII) biogenesis, activity, and assembly, together with coenzyme Q (CoQ) metabolism, are specifically impaired in the absence of SFXN1. Intriguingly, this cannot be explained by a defective 1C metabolism, which represents the only well-documented physiological role for SFXN1 (Kory et al., 2018). Instead, we show that SFXN1 is important for heme and central carbon metabolism, both of which support CIII function to varying extents. Collectively, these data define additional roles and thus provide a richer context for how SFXN1-mediated amino acid transport maintains mitochondrial integrity and promotes metabolic plasticity.

## RESULTS

### SFXN1 is a multipass inner mitochondrial membrane protein

SFXNs, consisting of 5 isoforms in humans (Figure S1A), are predicted to be integral polytopic IMM proteins with 4-5 TMDs (Fleming et al., 2001). The different isoforms have wide-ranging mRNA abundance across various tissues (Figure S1B). *SFXN1* is highly expressed in the kidney, small intestine, ovary, and especially the liver. *SFXN3* and *SFXN5* have notable expression in the brain, whereas *SFXN4* is the only isoform expressed well in heart and skeletal muscle. In HEK293, SFXN1 is the most abundant isoform based on protein level, followed by SFXN3 (Figure 1A) to which it shares the highest amino acid similarity (Figure S1A). SFXN1 is highly conserved across eukaryotic species and is closely related to yeast Fsf1p, having a 75% similarity and 37% identity (Figure S1C).

**Figure 1.**
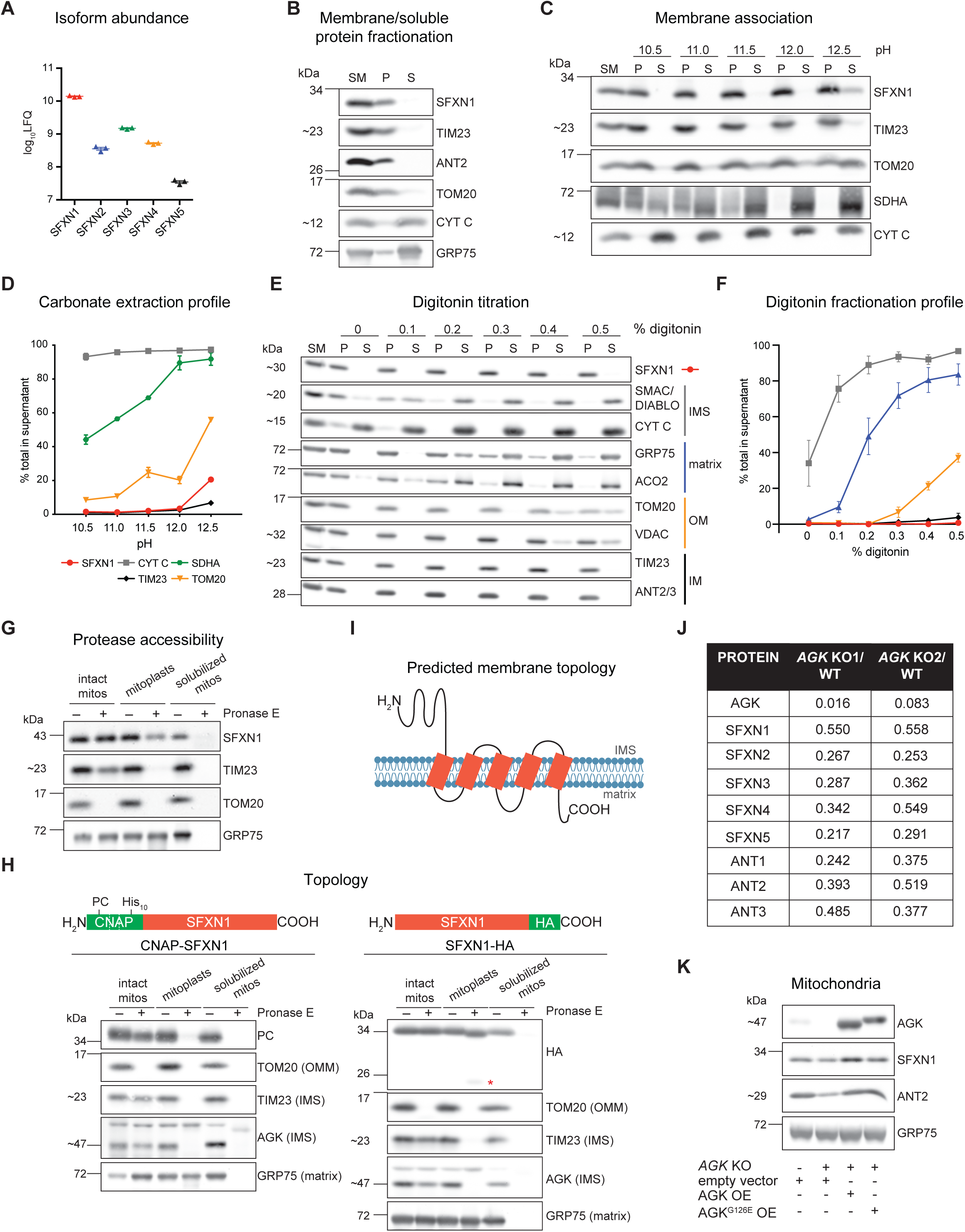
SFXN1, an integral inner mitochondrial membrane protein, is a TIM22 substrate. (A) Relative protein abundance of SFXN isoforms in HEK293 mitochondria as determined by mass spectrometry and label-free quantification (LFQ) (mean±SEM, *n*=3). (B) Sonication and centrifugation of mitochondria to separate membrane-bound from soluble proteins. SM, starting material; P, pellet; S, supernatant (C) Carbonate extraction of mitochondrial membrane proteins to distinguish between peripheral (appear in S) and integral (remain mostly in P) proteins. (D) Band intensities of P and S fractions in (C) were quantified and plotted as % of protein released in the supernatant (mean±SEM, *n*=3). (E) Digitonin titration for fractionation of mitochondrial subcompartments. Equal volumes of P and S fractions were analyzed. (F) Band intensities of P and S fractions (E) were quantified. Average band intensity of representative mitochondrial proteins from each subcompartment was plotted as % of protein released in the supernatant (mean±SEM, *n=3*). (G) Submitochondrial localization of endogenous SFXN1. HEK293 mitochondria were osmotically ruptured to yield mitoplasts, or solubilized with sodium deoxycholate. Samples were treated with Pronase E where indicated. (H) Submitochondrial localization of tagged SFXN1. HEK293 mitochondria lacking endogenous SFXN1 and expressing CNAP-SFXN1 or SFXN1-HA were processed as in (G). *, matrix-protected fragment (I) Predicted membrane topology of SFXN1 based on (H) (J) Proteomic analysis of *AGK* KOs versus WT. Shown are relative protein amounts of SFXN and ANT isoforms in the presence or absence of AGK. (K) Mitochondria from *AGK* KOs rescued with AGK, AGK^G126E^ or empty vector (v) were resolved by SDS-PAGE and immunoblotted for the indicated proteins.

SFXNs have been annotated as mitochondrial proteins based on large-scale proteomics (Calvo et al., 2016). We sought to biochemically characterize the membrane association, submitochondrial localization, and topology of SFXN1, the most abundant isoform in HEK293 (Figure 1A) (Lu et al., 2017). After sonication of mitochondria and ultracentrifugation to separate membrane-bound from soluble proteins, SFXN1 was found in the pellet (Figure 1B), meaning it is a membrane protein. To determine how it is associated with the membrane, we carried out carbonate extraction at increasing pH. In contrast to SDHA and CYT C, peripheral membrane proteins attached to membrane-spanning protein partners or the IMM, respectively, SFXN1 was mostly retained in the pellet and barely released from mitochondrial membranes even at pH 12.5 (Figure 1C). Its extraction profile is most similar to integral membrane proteins TIM23 and TOM20 (Figure 1D). Thus, SFXN1 possesses membrane-spanning domains.

We then investigated the submitochondrial localization of endogenous SFXN1 by digitonin titration (Figures 1E and 1F). Incubation of mitochondria with increasing digitonin concentrations results in the sequential release into the supernatant of IMS, matrix, outer mitochondrial membrane (OMM), and IMM proteins. The fractionation profile of SFXN1 resembled that of IMM residents. This result was further corroborated by a protease accessibility assay (Figure 1G). SFXN1 was only degraded when protease treatment was combined with detergent solubilization indicating that the epitope recognized by the SFXN1 antibody was not exposed to the IMS. This is in agreement with the localization of FLAG-tagged SFXN1 as determined by super-resolution microscopy (Kory et al., 2018).

Since there are conflicting results regarding the topology of SFXN1 (Lee et al., 2016; Lee et al., 2017), we performed a protease protection assay for N- and C-terminal epitope tagged SFXN1 (Figures 1H). After Pronase E treatment of mitoplasts, the CNAP tag of CNAP-SFXN1 was degraded similar to the IMS-exposed domains of TIM23 and AGK. However, Pronase E caused SFXN1-HA to migrate at a smaller size in mitoplasts, indicating that IMS-facing domain(s) in SFXN1 were degraded, and an even smaller band was detected that likely corresponds to an additional SFXN1-HA fragment that is matrix-protected. The HA tag was only degraded upon detergent inclusion. Altogether, these findings are in agreement with the study that mapped the topological direction of endogenous IMM proteins in live cells (Lee et al., 2017), and support that the N- and C-termini of SFXN1 face the IMS and the matrix, accordingly (Figure 1I). A model of SFXN1 with the two termini at opposing sides of the IMM is consistent with the notion that it possesses five TMDs and is thus likely to have a unique transport mechanism compared to SLC25 members.

### SFXNs are novel TIM22 substrates

MCs in the SLC25 family possess internal targeting elements and are integral polytopic proteins characterized by 6 TMDs. Their import and integration into the IMM is mediated by the TIM22 complex (Chacinska et al., 2009; Koehler, 2004; Neupert and Herrmann, 2007; Rehling et al., 2004). Recent studies have highlighted the substrate preference of TIM22 complex subunits and identified AGK as a metazoan-specific component essential for the import of MCs in the SLC25 family (Callegari et al., 2016; Kang et al., 2016; Kang et al., 2017; Vukotic et al., 2017). Since SFXN1 is a polytopic IMM protein, we asked whether SFXNs are also AGK-dependent TIM22 substrates. We employed SILAC-based proteomics coupled to LC-MS/MS on HEK293 (herein called wild type, WT) and *AGK* knockouts (KOs) generated by CRISPR/Cas9 (Figures S2A-S2C) (Mali et al., 2013; Ran et al., 2013), and found that protein levels of SFXN1-5, similar to ANT1-3, were reduced in the absence of AGK (Figure 1J). Given their dependence on AGK, the proteomics data also indicates that like SFXN1, the other SFXN isoforms are IMM residents in contrast to a recent report proposing that FLAG-tagged SFXN2 is in the OMM (Mon et al., 2019). As expected (Kang et al., 2017; Vukotic et al., 2017), SFXN1 protein abundance was not dependent on AGK kinase activity (Figure 1K and S2D; AGK^G126E^ is kinase-dead). In summary, SFXNs are novel TIM22 complex substrates whose steady-state abundance is dependent on AGK.

### Absence of SFXN1 does not severely impair iron homeostasis

To probe the physiological role of SFXN1, we generated *SFXN1* KOs in HEK293 by CRISPR/Cas9. We used two guide RNAs (gRNAs) targeting different parts of exon 2, and screened by immunoblotting (Figure S3A and S3B). Introduced genetic lesions in *SFXN1* KO1 and 2, generated by gRNA #1 and 2, respectively, were analyzed by Sanger sequencing. Both contained missense mutations leading to premature stop codons (Figure S3C).

SFXN1 was identified by positional cloning as the causative gene responsible for the phenotypic anomalies of the *flexed-tail* mouse, characterized by axial skeletal abnormalities, temporary embryonic and neonatal anemia, and mitochondrial iron deposits (Chui et al., 1977; Fleming et al., 2001; Hunt et al., 1933; Mixter and Hunt, 1933). However, other mutations potentially responsible for the observed deficiencies were subsequently found (Hegde et al., 2007; Lenox et al., 2005). Due to this, the role of SFXN1 in iron homeostasis has been debated. To test whether loss of SFXN1 disrupts iron homeostasis, we compared cell viability of WT and *SFXN1* KOs treated with increasing concentrations of deferoxamine mesylate (DFO). No difference between WT and *SFXN1* KOs was observed upon DFO-mediated iron chelation (Figure 2A) indicating that *SFXN1* KOs are not more sensitive to iron depletion and thus unlikely deficient in cellular iron. Indeed, cellular and mitochondrial iron levels as measured by ICP-MS were not altered in the absence of SFXN1 (Figures 2B and 2C). While the levels of other physiologically relevant metals were unaffected, cellular Mn^2+^ was decreased in *SFXN1* KOs. Since manganese plays a chief role as an enzyme cofactor, most notably for superoxide dismutases (Wang et al., 2018), this decrease in cellular Mn^2+^ could make *SFXN1* KOs more susceptible to oxidative stress. Collectively, these data provide evidence that the absence of SFXN1 alone does not impact iron homeostasis in HEK293.

**Figure 2.**
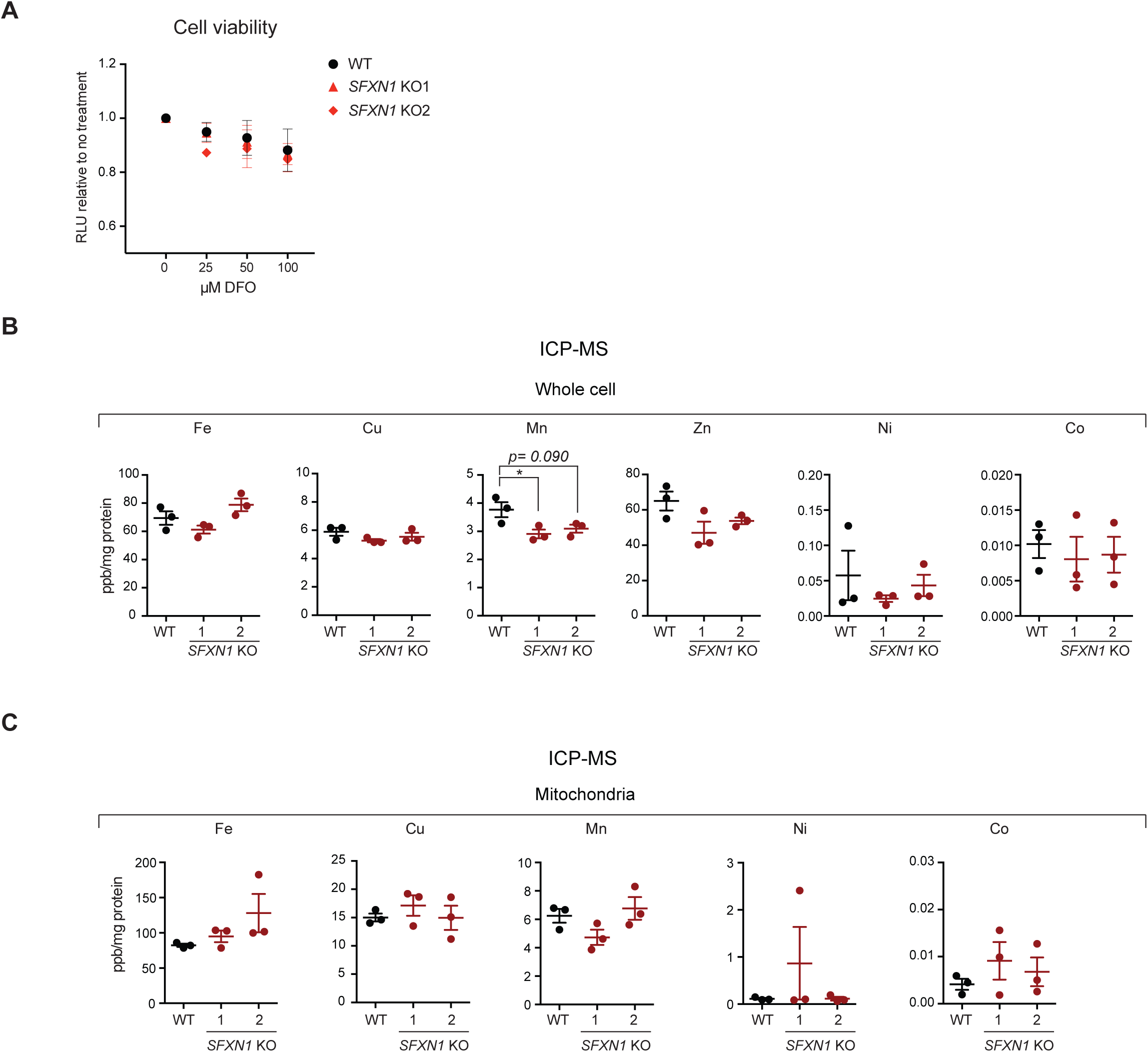
Deletion of *SFXN1* alone does not disrupt iron metabolism. (A) Cell viability of HEK293 WT and *SFXN1* KOs upon treatment with the indicated concentration of the iron chelator deferoxamine mesylate (DFO). (B) Relative metal concentrations in WT and *SFXN1* KO cells and (C) mitochondria as determined by ICP-MS (mean±SEM, *n=3*). *\*p<0.05, **p<0.01*, \*\*\**p<0.001*, unpaired Student’s t-test

### Loss of SFXN1 causes Complex III-related defects

Because no gross alterations in metal balance were observed in *SFXN1* KOs, we investigated the potential role of SFXN1 in general mitochondrial function. We analyzed the protein levels of respiratory complex subunits, import machinery components, and players important for mitochondrial morphology (Figures 3A, 3D and S3D). Comparing *SFXN1* KOs with WT, the level of UQCRC2, a Complex III (CIII) subunit, was significantly decreased in both KOs. Also, COX4 (COXIV) was slightly decreased and NDUFB6 (CI) was modestly increased. Other proteins involved in OXPHOS, biogenesis, or that reside in IMM or matrix were unaffected or inconsistently altered in KOs relative to WT. A limited survey of electron transport chain (ETC) subunits containing iron-sulfur (Fe-S) clusters determined that in *SFXN1* KOs, UQCRFS1 (CIII) was remarkably decreased, SDHB (CII) was significantly reduced, and NDUFS1 (CI) was unaltered (Figures 3B and 3D), indicating that SFXN1 absence impacts a subset of Fe-S cluster-containing mitochondrial proteins. Since the tested CIII subunits were reduced upon SFXN1 loss, we asked whether the protein abundance of MTCYB, the sole mtDNA-encoded CIII subunit, was also altered. In *SFXN1* KOs, MTCYB protein, which was barely detectable in WT cells treated with doxycycline (dox) to inhibit mitochondrial translation, was reduced by ∼75% relative to untreated WT (Figures 3C and 3D). Altogether, these data provide evidence that *SFXN1* deletion is most detrimental to CIII subunit protein levels.

**Figure 3.**
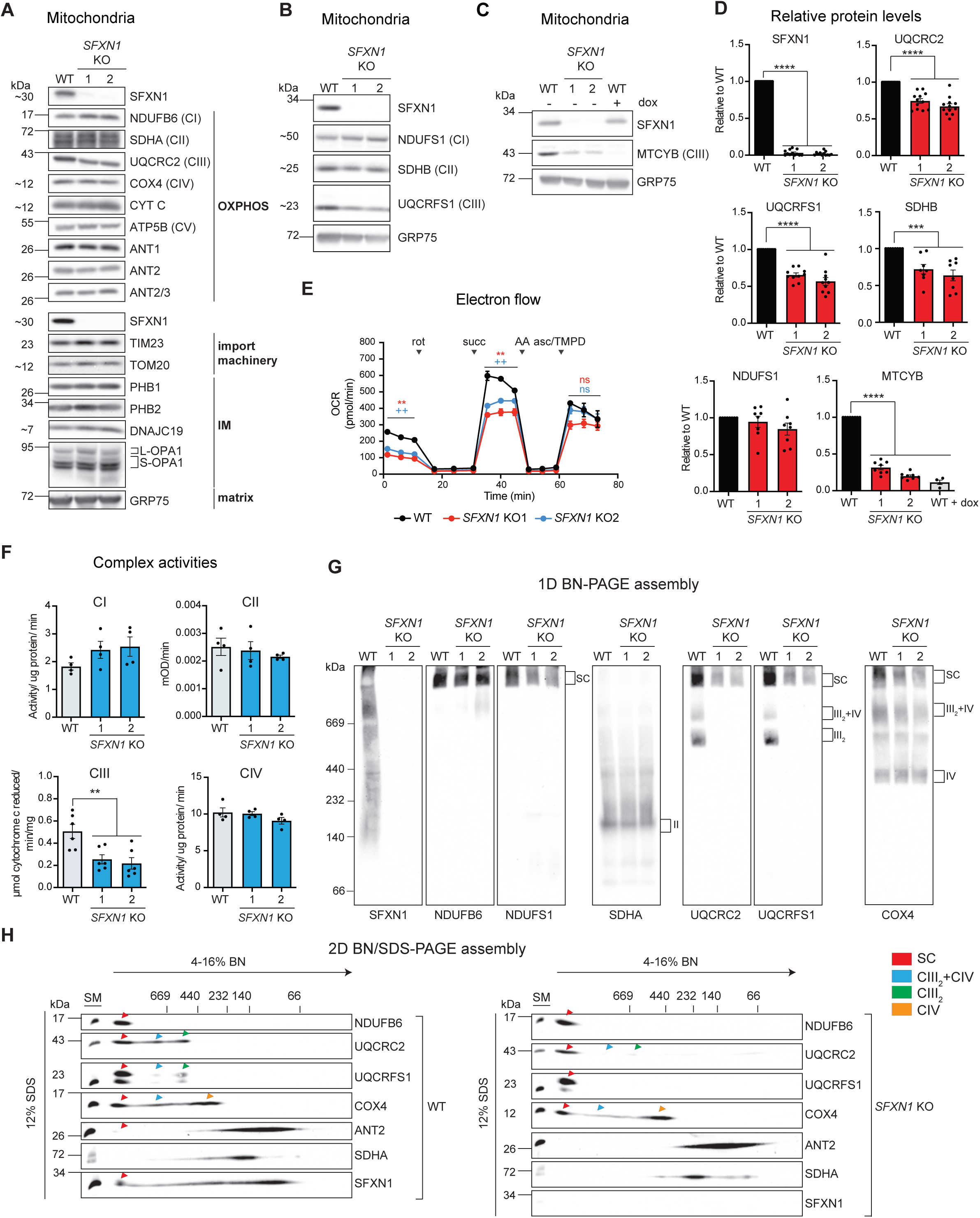
Absence of SFXN1 leads to Complex III-related defects. (A) Steady-state abundance of select mitochondrial proteins including OXPHOS components, subunits of import machineries, and other proteins in the IMM and matrix. (B) Immunoblotting for Fe-S containing subunits of respiratory complexes in mitochondrial extracts. GRP75 served as loading control. (C) Immunoblotting for MTCYB using mitochondrial extracts. Doxycycline-treated WT (+dox) shows that MTCYB was reduced when mitochondrial translation is halted. (D) Densitometric analysis of bands for select proteins in (A), (B) and (C). Protein levels in WT were set to 1.0 (mean±SEM, *n*≥4). (E) Electron flow through the respiratory chain in intact mitochondria. Base buffer contains 10 mM pyruvate, 2 mM malate and 4 µM CCCP. Injections of complex inhibitors/substrates were performed as indicated. rot, rotenone; succ, succinate; AA, antimycin A; asc/TMPD, ascorbate/TMPD. (F) Spectrophotometric assays using detergent-solubilized mitochondria to monitor individual complex activities. CI, oxidation of NADH to NAD^+^; CII, ubiquinol production; CIII-cytochrome c reduction; CIV-cytochrome c oxidation (mean±SEM, *n*≥4). (G) 1D BN assembly. Mitochondria solubilized in 1% (w/v) digitonin were resolved on a 4-16% BN gel and immunoblotted for the indicated subunits. (H) Mitochondria solubilized with 1% (w/v) digitonin were resolved via 2D BN/SDS-PAGE and immunoblotted for the indicated subunits. \**p<0.05*, \*\**p<0.01*, \*\*\**p<0.001*, unpaired Student’s t-test

To determine whether *SFXN1* ablation leads to non-optimal ETC function, we assessed electron flow through ETC complexes by sequentially feeding specific substrates and inhibitors to intact mitochondria (Figure 3E). *SFXN1* KO mitochondria registered significantly lower oxygen consumption rates (OCRs) when pyruvate (CI substrate) or succinate (CII substrate) was supplied, but not after addition of ascorbate/TMPD (CIV substrate), indicating that there is a problem pre-CIV. To further isolate the pre-CIV ETC defect, the individual CI to CIV activities were measured in detergent-solubilized mitochondria which demonstrated a specific defect in CIII (Figure 3F and Figure S3E).

Next, we interrogated the effect of *SFXN1* deletion on the capacity of respiratory complexes to associate into higher order structures. Digitonin-solubilized mitochondria were subjected to 1D Blue Native (BN) (Figure 3G) and 2D-BN/SDS-PAGE (Figure 3H). The absence of SFXN1 did not prevent the formation of respiratory supercomplexes (SC). Strikingly, in *SFXN1* KOs, the CIII_2_-CIV subcomplex and CIII_2_ were prominently decreased when immunoblots were probed with antibodies against two CIII subunits (UQCRC2 and UQCRFS1). Using the COX4 antibody, this reduction in CIII_2_-CIV subcomplex, which was more obvious in 2D BN/SDS-PAGE, was confirmed. CI assembly, monitored using NDUFB6 and NDUFS1 antibodies, appeared normal. Similarly, ANT2 and SDHA assemblies were unchanged in the absence of SFXN1 (Figure 3H). The reduction in CIII_2_-CIV was still apparent even following extraction with DDM, a detergent that unlike digitonin, destabilizes many of the larger SCs (Figure S3F). Moreover, there were no changes in the overall electrophoretic pattern of complexes analyzed by 1D-BN and Coomassie staining (Figure S3G). Knocking-out *SFXN1* in HeLa also rendered CIII dysfunctional (Figures S4A-S4C). Moreover, stably overexpressing SFXN1 in *SFXN1* KO partially improved CIII function (Figures S4D and S4E). We speculate that reintroducing SFXN1 at a physiological level may provide a better rescue as overexpression of MCs has been shown to induce an aggregation-based, cytosolic stress response (Liu et al., 2019).

In *SFXN1* KOs, the lower CIII subunit protein levels and activity, and the specific reduction of CIII_2_-CIV and CIII_2,_ cannot be accounted for by a transcriptional mechanism (Figure S4F), or by changes in mtDNA content (Figure S4G). They also cannot be readily explained by a mitochondrial translation defect since only a modest impairment in MTCYB translation was noted (Figures S4H and S4I).

CIII couples electron transfer from CoQ to CYT C to proton pumping into the IMS. The mammalian CIII monomer is composed of 11 subunits. Assembly of dimeric CIII begins with the incorporation of chaperone-bound MTCYB into the IMM followed by the attachment of nuclear-encoded structural subunits to form pre-CIII_2_ (Fernandez-Vizarra and Zeviani, 2018; Signes and Fernandez-Vizarra, 2018). BCS1L and LYRM7 are assembly factors required for the proper incorporation of UQCRFS1 into pre-CIII_2_ and thus the full maturation of active CIII_2_ (Fernandez-Vizarra and Zeviani, 2015). Notably, the CIII assembly defect observed in *SFXN1* KOs is reminiscent of that documented in BCS1L-mutant and LYRM7^HA^-overexpressing cell lines (Bottani et al., 2017; Fernandez-Vizarra et al., 2007; Sanchez et al., 2013). However, BCS1L and LYRM7 protein levels were not altered in the absence of SFXN1 (Figure S4J). These results suggest that SFXN1 influences CIII integrity by another mechanism.

Based on this possibility, we reasoned that impaired CIII function may affect the oxidation of CoQ, a lipid that transfers electrons from CI/CII to CIII, which in turn can impact CoQ content. We therefore measured its two isoforms in humans: CoQ_9_ and CoQ_10_. Both were markedly reduced in the absence of SFXN1, and total CoQ in *SFXN1* KOs was ∼50% that of WT (Figure 4A). Since saturating amount of reduced CoQ was utilized in CIII activity assays (Figures 3F and S3E) and CYT C protein levels were largely unaffected (Figures 3A and S3D), SFXN1-null cells are characterized by an intrinsic CIII defect that may affect CoQ biosynthesis and/or turnover.

**Figure 4.**
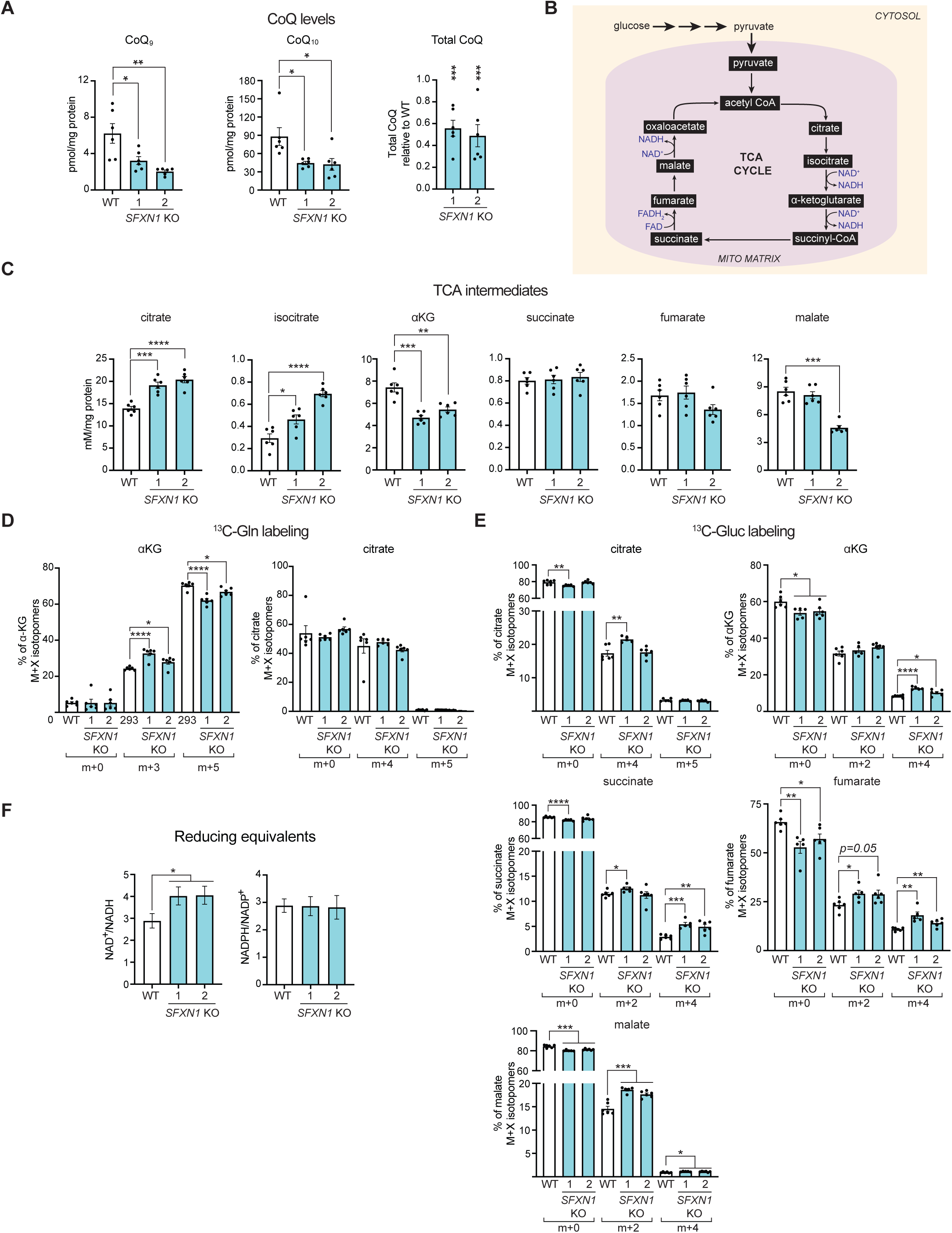
SFXN1 loss perturbs CoQ and central carbon metabolism. (A) CoQ measurements in cellular lipid isolates by RP-HPLC/MS (mean±SEM, *n=6*). (B) TCA cycle (C) Steady-state intracellular concentrations of TCA cycle metabolites obtained by LC-MS/MS (mean±SEM, *n=6*). αKG, α-ketoglutarate. (D,E) Percent abundance of m+x-labeled metabolites as determined by LC-MS/MS upon (D) [U-^13^C]-glutamine labeling (mean±SEM, *n=6*) and (E) [U-^13^C]-glucose labeling (mean±SEM, *n*≥*5*). (F) Quantification of NAD^+^/NADH (mean±SEM, *n*≥*8*) and NADPH/NADP^+^ ratios (mean±SEM, *n=6*). \**p<0.05*, \*\**p<0.01*, \*\*\**p<0.001*, unpaired Student’s t-test

### *SFXN1* deletion affects central carbon metabolism

SFXN1 transports serine and, based on their ability to inhibit serine flux across the IMM, additional physiologically important amino acids including alanine, cysteine and glycine (Kory et al., 2018). Thus, we explored other metabolic consequences upon SFXN1 loss. Amino acid metabolism is tightly coupled with the tricarboxylic acid (TCA) cycle. As such, the steady-state levels of TCA intermediates were determined (Figure 4B). In *SFXN1* KOs, citrate and isocitrate were elevated while α-ketoglutarate (αKG) was reduced (Figure 4C), suggesting that SFXN1 is important for central carbon metabolism. To better understand these changes, WT and *SFXN1* KOs were incubated with [U-^13^C]-glutamine and isotopomer distribution into TCA metabolites examined (Figures 4D, S5A-S5B). *SFXN1* KOs had higher m+3-labeled αKG representing acetyl-CoA contribution to αKG but lower m+5-labeled αKG, indicating inefficient conversion of glutamine to αKG. This is consistent with changes in activities of enzymes involved in αKG metabolism in the absence of SFXN1 (Figures S5D-S5G). Activity of GDH, which mediates αKG formation from glutamate (Hudson and Daniel, 1993), was decreased in *SFXN1* KOs. Activity of ALT, which facilitates the reversible addition of an amino group from L-alanine to αKG to yield pyruvate and glutamate (Groen et al., 1982; Yang et al., 2002), was also decreased in *SFXN1* KOs, likely due to the lower αKG in these samples. The elevated citrate/isocitrate in *SFXN1* KOs was not due to changes in oxidative (m+4-labeled citrate) or reductive glutamine utilization (m+5-labeled citrate) (Figure 4D).

[U-^13^C]-glucose tracing (Figures 4E and S5C) suggests that in the absence of SFXN1, there is an increase in glucose incorporation into TCA intermediates particularly after the first round of the cycle (m+4). Also, we observed a higher NAD^+^/NADH with normal NADPH/NADP^+^ ratio in *SFXN1* KOs (Figure 4F), consistent with a disturbance in central carbon metabolism.

Despite these metabolic alterations, mitochondrial respiration (Figure 5A) and glycolytic parameters (Figure 5B) were not impacted in intact *SFXN1* KO cells. This is in stark contrast to isolated mitochondria, where the loss of SFXN1 diminished basal and/or ADP-stimulated oxygen consumption driven by various fuels that donate electrons to CI, CII and CIII, in agreement with a CIII defect (Figure 5C). *SFXN1* KO mitochondria also displayed a lower capacity to oxidize palmitoylcarnitine/malate, suggesting a faulty coordination between β-oxidation and the ETC. We therefore asked how *SFXN1* KOs are able to maintain cellular baseline respiration, and observed higher dependency on glucose oxidation, with largely unaltered reliance on glutamine and fatty acid pathways (Figure 5D), consistent with the increase in glucose contribution to TCA metabolites (Figure 4E). Overall, this highlights the metabolic plasticity at the cellular level driven by the loss of SFXN1.

**Figure 5.**
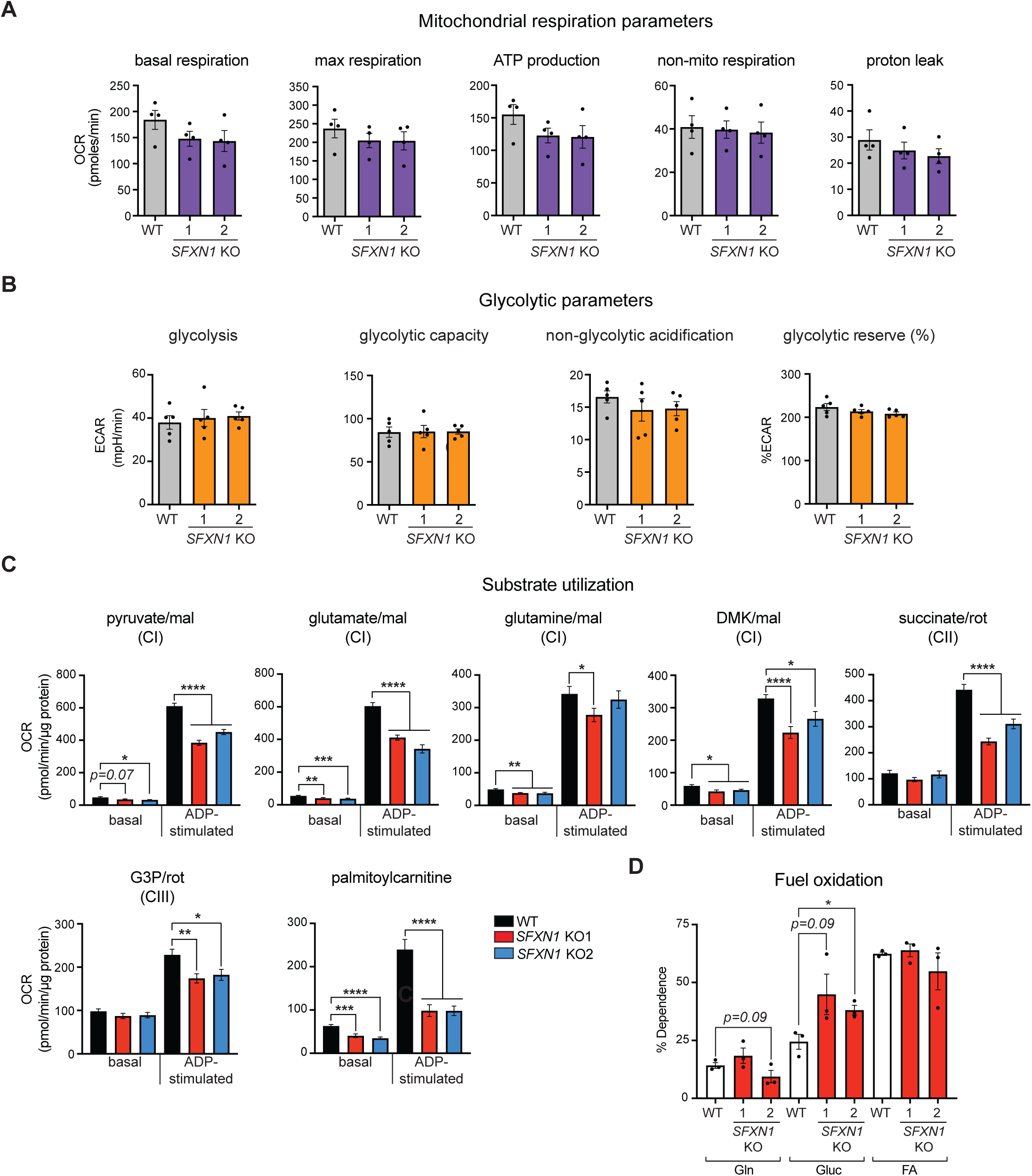
Cells lacking SFXN1 display metabolic flexibility. (A) Oxygen consumption rate (OCR) in intact cells. Presented are values normalized by DNA content (mean±SEM, *n=4*). (B) Extracellular acidification rate (ECAR) in intact cells. Presented are values normalized by DNA content (mean±SEM, *n=5*). (C) OCR in isolated mitochondria using the specified substrates (mean±SEM, *n=7*). mal, malate; rot, rotenone; G3P, glycerol-3-phosphate (D) Percent dependency on glutamine (Gln), glucose (Gluc), and fatty acid (FA) oxidation (mean±SEM, *n=3*). \**p<0.05*, \*\**p<0.01*, \*\*\**p<0.001*, unpaired Student’s t-test

### *SFXN1* deletion impedes heme biosynthesis

Next, we assessed the relative amounts of two substrates of SFXN1 (Kory et al., 2018). Total serine in *SFXN1* KOs was elevated while total glycine was unmodified in comparison to WT (Figure S5H). Previous work reported that SFXN1 loss in Jurkat and K562 cells leads to increased serine and decreased glycine levels (Kory et al., 2018). Despite the difference in the effect on glycine, both studies identify a higher serine:glycine ratio in cells lacking SFXN1. Thus, the absence of SFXN1 results in serine accumulation potentially due to inefficient serine to glycine conversion.

Serine reaching the mitochondrial matrix has several metabolic fates (Mattaini et al., 2016). Its cleavage yields 5,10-methylenetetrahydrofolate (5,10-methyl-THF) and glycine. 5,10-methyl-THF feeds into 1C metabolism that contributes to nucleotide biosynthesis (Lewis et al., 2014) while glycine serves as a precursor for glutathione (Wu et al., 2004) and heme (Radin et al., 1950). SFXN1 has been shown to play a role in 1C metabolism (Kory et al., 2018), but whether it affects other branches of serine catabolism vital for mitochondrial function has not been addressed. We hypothesized that insufficient serine and/or glycine import into mitochondria upon SFXN1 loss might also impair heme generation, a tightly regulated, multi-step metabolic pathway (Figure 6A). Consistent with this possibility, the mRNA level of *CPOX* and *FECH* (Figure 6B) were reduced in the absence of SFXN1. Steady-state protein amount of FECH (Figure 6C), an Fe-S containing protein that catalyzes Fe^2+^ insertion into protoporphyrin IX (PPIX) (Ajioka et al., 2006), was also decreased in the absence of SFXN1 and could be partially rescued upon its re-introduction (Figure S6B). ALAS1, which performs the initial and rate-limiting step of heme production (Figure 6A), was increased in *SFXN1* KOs compared to WT (Figure 6C). *ALAS1* is subject to negative feedback-mediated regulation by heme (Kubota et al., 2016), suggesting that heme levels may be limiting when SFXN1 is missing. Indeed, heme was reduced in *SFXN1* KOs (Figure 6D), consistent with a role for SFXN1 in promoting heme biosynthesis.

**Figure 6.**
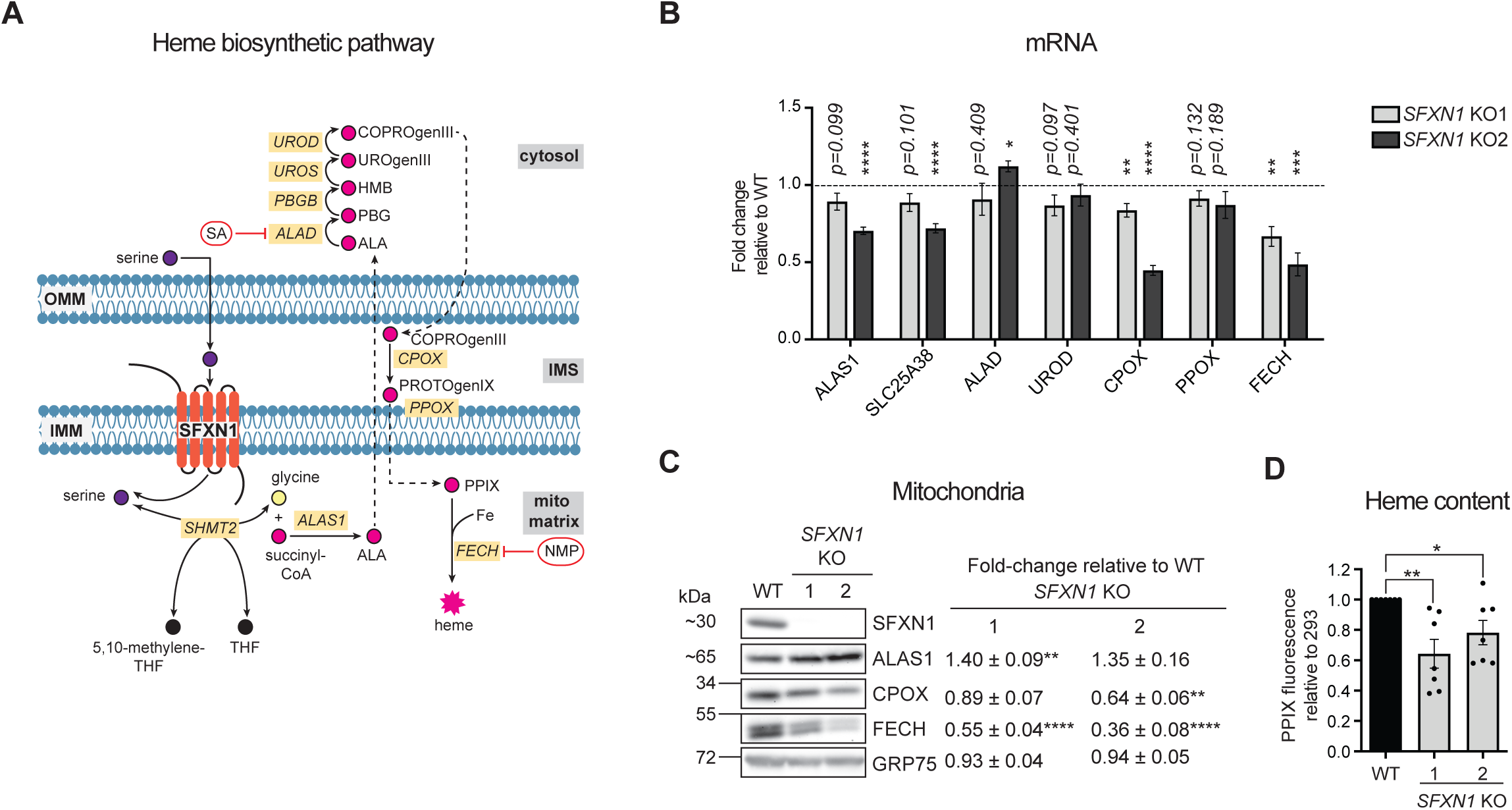
Heme biosynthesis is compromised in *SFXN1* KOs. (A) Heme biosynthetic pathway. SA, succinylacetone; NMP, N-methyl protoporphyrin; PPIX, protoporphyrin IX (B) Gene expression analysis of heme biosynthetic enzymes by qPCR (mean FC±SEM, *n=4*). (C) Immunoblotting for enzymes involved in heme biosynthesis. Values shown are fold-change protein steady-state abundance in *SFXN*1 KOs over HEK293 WT (mean±SEM, *n=5*). (D) Relative cellular heme levels obtained by analyzing PPIX fluorescence (mean±SEM, *n=7*) \**p<0.05*, \*\**p<0.01*, \*\*\**p<0.001*, unpaired Student’s t-test

### Hemin, but not formate, supplementation improves CIII function in *SFXN1* KOs

Because SFXN1 has the ability to transport several amino acids *in vitro* (Kory et al., 2018), we tested whether inefficient transport of a particular substrate into the mitochondrion can explain the CIII defects observed in *SFXN1* KOs. Exogenous serine, glycine, alanine or cystine did not restore CIII subunit protein abundance in *SFXN1* KOs to WT levels, and had no obvious effect on other ETC proteins assessed (Figure S5I). It remains a possibility that an amino acid cocktail instead of an individual amino acid can account for the defects observed. It is also plausible that without SFXN1, the specific amino acid supplied cannot robustly reach the mitochondrial matrix to exert an effect. If true, this would imply that other MCs, including other SFXNs, may be partially, but not completely, functionally redundant with SFXN1. We therefore asked whether suboptimal generation of specific metabolites in the mitochondrial matrix due to inefficient transport of precursors could better explain the CIII-related phenotype of *SFXN1* KOs. Two candidate metabolites are formate and heme. Formate, one of the most studied products of serine catabolism, is involved in mitochondrial translation and thus can affect respiratory chain function (Lucas et al., 2018; Minton et al., 2018; Morscher et al., 2018). On the other hand, heme moieties are present in CIII (Kim et al., 2012), thus it is possible that the CIII dysfunction in *SFXN1* KOs stems from their impaired heme metabolism (Figure 6).

In cells lacking SFXN1, formate shortage is responsible for their slower proliferation in serine-deficient media (Kory et al., 2018). We noted WT already displayed a proliferation defect upon serine withdrawal in glucose-based media, implying that *de novo* serine synthesis is not sufficient for normal cell proliferation in HEK293 cells (Figure 7A). This was exacerbated when *SFXN1* was deleted. Formate supplementation, but not hemin, a cell-permeable derivative of heme (Nakamichi et al., 2005; Wang et al., 2009), rescued this phenotype.

**Figure 7.**
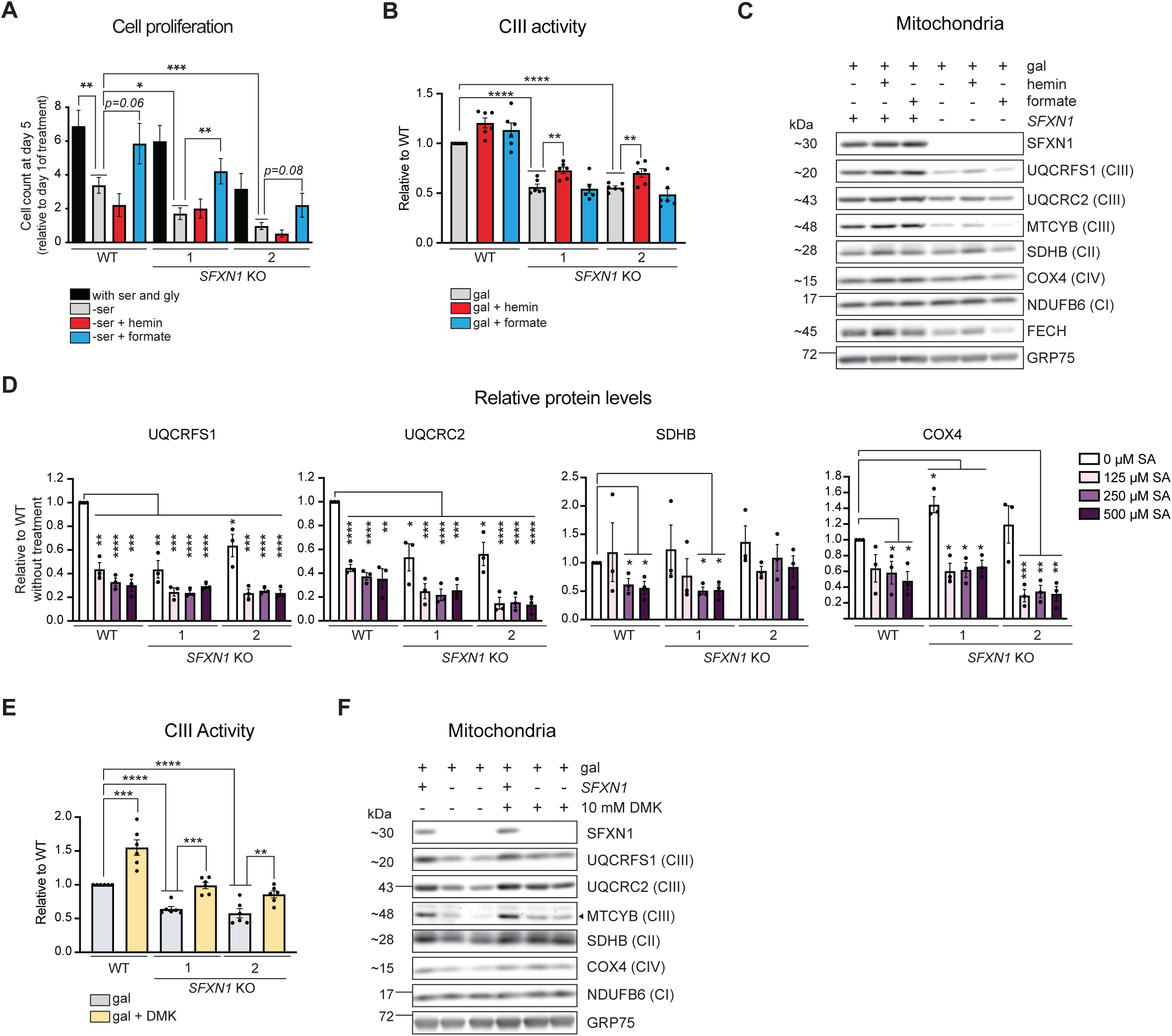
Reinforcement of heme and central carbon metabolism, but not the one-carbon pathway, partially restores Complex III function in the absence of SFXN1. (A) Cell proliferation in full media (with serine), without serine (-ser), and upon supplementation of serine-free media with 15 µM hemin (-ser + hemin) or 1 mM formate (-ser + formate) (mean±SEM, *n≥4*). (B) Complex III activity in DDM-solubilized mitochondria. 48 hours before mitochondrial isolation, cells were switched to media containing galactose only, galactose with 15 μM hemin or galactose with 1 mM formate (mean±SEM, *n=6*). (C) Immunoblotting for select respiratory complex subunits using mitochondrial isolates detailed in (B). (D) Steady-state protein abundance of select respiratory complex subunits in cell lysates. Cells were grown in glucose-based media and treated with the indicated concentration of succinylacetone (SA), an inhibitor of heme biosynthesis for 2 days. (E) Complex III activity in DDM-solubilized mitochondria. 48 hours before mitochondrial isolation, cells were switched to media containing galactose only, or galactose with 10 mM dimethyl α-ketoglutarate (DMK) (mean±SEM, *n=6*). (F) Immunoblotting for select respiratory complex subunits using mitochondrial isolates detailed in (E). \**p<0.05*, \*\**p<0.01*, \*\*\**p<0.001*, unpaired Student’s t-test

Curiously, other studies have determined that formate depletion mainly affects activities of CI, CIV and CV and their subunit protein abundances, but not CIII (Lucas et al., 2018; Minton et al., 2018; Morscher et al., 2018). Indeed, exogenous formate did not reverse the reduced CIII activity (Figure 7B) and CIII subunit protein levels (Figure 7C) in *SFXN1* KOs. The inability of formate to restore CIII expression or activity is consistent with the preserved mtDNA translation in *SFXN1* KOs (Figures S4H and S4I). In contrast to formate, exogenous hemin enhanced CIII activity in both WT and *SFXN1* KOs (Figure 7B). These results indicate that in absence of SFXN1, the 1C pool is a limiting factor for nucleotide synthesis and therefore cell proliferation (Kory et al., 2018; Labuschagne et al., 2014) whereas deficient heme is partially responsible for the decreased CIII function.

We noted that serine withdrawal from glucose-containing media is not sufficient to impact CIII activity or CIII subunit protein levels in WT or *SFXN1* KOs (Figures S6A and S6C). Still, CIII impairments manifested in *SFXN1* KOs even with glucose as carbon source instead of galactose, wherein cells were not forced to respire (Figures S6B, S6C and S6D). In this metabolic setting, supplementation with hemin, but not formate, again improved CIII activity in cells lacking SFXN1 (Figure S6B).

Addition of hemin, however, only minimally affected CIII subunit protein abundance and assembly in *SFXN1* KOs regardless of carbon source (Figures 7C, S6C and S6D). This suggests that the lower heme resulting from SFXN1 loss contributes to the decreased CIII activity, but that there are other metabolites formed following SFXN1-mediated transport of serine or other amino acids that can influence CIII stability. It also remains a possibility that exogenous hemin is not as readily utilized for processes that ensure proper CIII expression and assembly.

To interrogate whether heme deficiency affects protein abundance of ETC complex subunits independent of a perturbed serine metabolism, we treated cells with increasing concentrations of succinylacetone (SA), which blocks heme biosynthesis by inhibiting ALAD (Figure 6A) (Sassa and Kappas, 1983; Tschudy et al., 1981). SA treatment decreased protein levels of CII, CIII and CIV, but not CI which does not bear heme, and upregulated ALAS1 in response to the expected aminolevulinic acid accumulation (Maitra et al., 2019) (Figures 7D, S7A and S7B). While CII (SDHB) and CIV (COX4) protein levels were reduced by ∼50% upon treatment of WT cells with 500 µM SA, CIII (UQCRC2 and UQCRFS1) was decreased by ∼75% (Figures 7D and S7A). This suggests that CIII expression and/or stability is very sensitive to SA-mediated heme deprivation. Deleting SFXN1 furthered this drop in CIII subunit protein abundance upon SA treatment (Figures 7D and S7A). This is in agreement with the already reduced heme levels in these cells, and also indicative that other roles of SFXN1 contribute to the CIII-related phenotype. On the other hand, treatment with N-methyl protoporphyrin (NMP), which represses FECH by being a PPIX transition analog (Figure 6A), only affected CIV (Figure S7C). Selective susceptibility of CIV assembly and subunit protein levels to NMP treatment has been previously documented (Atamna et al., 2001). Based on our data, this seems to be a NMP-specific response, suggesting that this drug may preferentially affect heme *a* that is only present in CIV (Atamna et al., 2001; Kim et al., 2012). In summary, inadequate heme may also be a contributing factor to the lower CIII subunit protein levels in *SFXN1* KOs.

### DMK supplementation restores CIII upon *SFXN1* ablation

Because αKG was reduced in *SFXN1* KOs, we asked whether addition of dimethyl α-ketoglutarate (DMK), a cell-permeable version of αKG, has a positive effect on CIII function. Surprisingly, DMK supplementation fully rescued CIII activity in *SFXN1* KOs to WT level and even enhanced WT CIII activity by ∼50% (Figure 7E). It also rescued CIII subunit protein abundance in *SFXN1* KOs, and increased the levels of SDHB and COX4, which were also slightly down in cells lacking SFXN1 (Figure 7F). In toto, our data support a model in which SFXN1 regulates CIII expression and activity by both promoting heme synthesis and maintaining an intact central carbon metabolism, possibly by fine-tuning flux into the TCA cycle.

## DISCUSSION

MCs represent important nodes of metabolism. In this study, we uncover additional functions for SFXN1, an unconventional MC that transports select amino acids including serine, outside of its role in the generation of 1C units (Kory et al., 2018). We demonstrate that SFXN1 is important for CIII integrity, as well as CoQ, heme and central carbon metabolism.

With a series of biochemical experiments, we show that SFXN1 is an integral membrane protein in the IMM with N- and C-termini exposed to the IMS and mitochondrial matrix, respectively, consistent with having five TMDs. Also by a protease protection assay, another study has shown that the N-terminus of FLAG-SFXN4 is facing the IMS (Hildick-Smith et al., 2013). Together with SFXN1, SFXN4 and SFXN2 are modeled to have five TMDs, while SFXN3 and SFXN5 are predicted to have only four. Whether or not this feature imparts substrate preference of each isoform is unknown. Structural information of the SFXN family, including insight into determinants of ligand selectivity that may dictate the transport mechanism, are all currently lacking.

Based on proteomics data, we show that together with SLC25 family members including the prototypical ANTs (Kang et al., 2017; Vukotic et al., 2017), SFXNs are AGK-reliant TIM22 cargos. Interestingly, there seems to be a range of AGK-dependency for the different ANT and SFXN isoforms (Figures 1J and 1K). This suggests that there are perhaps unknown players required for the import/assembly of specific carriers, or that the number of TMDs or some other structural element may govern their dependence on AGK. Orthodox TIM22 substrates contain 4 or 6 TMDs (Rehling et al., 2003), but a recent report using yeast, which lacks AGK, expanded this catalog to include subunits of the mitochondrial pyruvate carrier, consisting of 2 or 3 TMDs (Gomkale et al., 2020). The odd-number of TMDs in SFXN1 thus makes it another atypical TIM22 substrate.

We find that SFXN1 is vital for CIII integrity and CoQ metabolism. Being a mitochondrial serine transporter, SFXN1 is positioned to influence numerous mitochondrial functions, as there is a close relationship between ETC efficiency and serine metabolism. It has been shown that mtDNA depletion activates serine biosynthesis, and that respiratory complex lesions perturb 1C metabolism by limiting formate production from serine (Bao et al., 2016; Nikkanen et al., 2016). Our study contributes to growing evidence that conversely, disruption of serine metabolism also exerts a toll on mitochondrial function. Supplementation of formate, which maintains tRNA methylation needed for proper mitochondrial protein translation (Lucas et al., 2018; Minton et al., 2018; Morscher et al., 2018), did not restore any tested aspect of CIII function. This indicates that a problematic 1C metabolism is not the underlying cause of any of the SFXN1-related CIII defects as defined here. Serine deprivation has been shown to induce changes in lipid profile that subsequently leads to mitochondrial fragmentation (Gao et al., 2018), serving as precedent that serine-derived metabolites other than 1C units influence mitochondrial physiology. These findings also argue that SFXN1 is more than just a player in 1C metabolism.

We illustrate that in the context of CIII dysfunction, the inefficient generation of at least two metabolites —heme and αKG— contributes to the phenotype. SFXN1 is linked to heme biosynthesis by its ability to deliver serine, and perhaps glycine (Kory et al., 2018), into the mitochondrial matrix where the pathway is initiated. Consistently, SFXN1 is a constituent of the mitochondrial heme metabolon (Medlock et al., 2015). In endothelial cells, depletion of cellular serine by deletion of *PHGDH*, encoding the rate-limiting enzyme in *de novo* serine biosynthesis, results in inadequate heme and impedes OXPHOS capacity (Vandekeere et al., 2018). In our HEK293-based *SFXN1* KO model, there is an accumulation of total serine and an elevated serine:glycine ratio, implying that it is the ability to take serine into the mitochondrion for glycine conversion and not low serine levels per se that underlies the observed heme-related defect. Interestingly, although accumulation of serine in the cytosol may in theory increase its rate of conversion to glycine that in turn may possibly be transported to the mitochondrion (Guernsey et al., 2009; Lunetti et al., 2016), it seems that this is insufficient to maintain normal heme levels. This highlights the significance of achieving threshold metabolite concentrations in specific subcellular compartments to drive biosynthetic processes.

Notably, other SFXNs have been implicated in heme metabolism, including SFXN4 which has not been demonstrated to compensate for serine transport-based derangements in the absence of SFXN1 (Kory et al., 2018; Mon et al., 2019; Paul et al., 2019). At least in murine erythroleukemia cells, glutamine is the major source of succinyl-CoA for aminolevulinic acid (ALA) production (Burch et al., 2018). It is thus conceivable that SFXN4 and perhaps other isoforms can impact heme generation by transporting amino acids other than serine.

Consistent with a contribution of reduced heme content to the CIII-associated phenotype, we show that hemin supplementation improved CIII activity in cells lacking SFXN1. This is in agreement with other studies showing that heme deprivation is associated with lower activities of heme-containing RCs II-IV (Gamble et al., 2000; Lin et al., 2019; Vandekeere et al., 2018). CIII subunit protein levels and assembly proved to be more difficult to rescue with hemin. Potentially there is a difference in the bioavailability of exogenously supplied hemin and endogenously synthesized heme for incorporation into CIII that subsequently affects its stability. Suboptimal delivery of exogenous heme to mitochondria has been previously documented (Kim et al., 2016). This may be a potential explanation why hemin supplementation usually results in only a partial rescue of RC function (Shetty et al., 2019; Vandekeere et al., 2018).

We showed that impeding heme production at two unique steps of the pathway differentially affects RC subunits. SA, which acts at an early point by inhibiting ALAD, reduces steady-state protein levels of CII-CIV, with CIII subunits being most sensitive. However, NMP, which blocks the last step by inhibiting FECH, only decreased CIV. Interestingly, although both are known to deplete heme, we observed that SA and NMP treatments have opposite effects on ALAS1 protein (Figures 7D, S13A and S14A). SA caused upregulation, possibly as a response to ALA buildup, while NMP led to downregulation, likely due to PPIX accumulation (Maitra et al., 2019). Hence, the impact of heme metabolism on RC subunit protein levels and assembly appears to be more complex than being just about total heme amount. Going forward, it will be important to consider how genetic or pharmacological interventions affect heme biosynthetic intermediates and specific heme types.

*SFXN1* KOs displayed imbalanced central carbon metabolism as indicated by a substantial reduction in αKG, in part due to ineffective glutamate conversion to αKG, and elevation in citrate/isocitrate. Supplying αKG to the TCA cycle is relevant for the production of NADH and FADH_2_. αKG is connected to the serine synthesis pathway— glutamate, acting as a nitrogen donor, is converted to αKG as phosphoserine is formed from PHGDH-generated 3-phosphohydroxypyruvate (Amelio et al., 2014). It will be compelling to obtain a quantitative perspective of other metabolic flux alterations in the absence of SFXN1. The reversal of CIII activity and subunit protein abundance in *SFXN1* KOs upon DMK treatment emphasizes the tight coordination of flux into the TCA cycle and OXPHOS. In cell models with moderate OXPHOS decay, stimulation of αKG anaplerosis by exogenous DMK is beneficial for cell proliferation and respiration in galactose-based media (Chen et al., 2018). Here we provide evidence that these may be mediated by elevation in CIII activity and protein levels of RCs. Exactly how αKG does this warrants further investigation.

Why is CIII more sensitive to *SFXN1* ablation than other RCs? Although there is presently no clear answer to this enigma, it is intriguing that SFXN1 has been documented as a binding partner of TTC19, a CIII assembly factor (Bottani et al., 2017). It is becoming increasingly appreciated that different aspects of mitochondrial biology are subject to metabolic regulation, from mitochondrial biogenesis (Scarpulla, 2011) to dynamics (Mishra and Chan, 2016). An emerging form of metabolic communication is mediated by metabolite-dependent post-translational modifications of proteins that modulate ETC biogenesis, assembly and activation (Van Vranken et al., 2018). It is an interesting possibility that MCs, by delivering substrates, work closely with these factors.

Overall, our functional investigation significantly extends our understanding of how SFXN1 influences mitochondrial and cellular metabolic efficiency. In light of the data indicating that SFXN1 likely transports other amino acids besides serine (Kory et al., 2018), together with the documented substrate promiscuity of MCs (Boulet et al., 2018; Fiermonte et al., 2009; Seifert et al., 2015; Wohlrab and Flowers, 1982), it is plausible that SFXN1 participates in other anabolic and catabolic reactions beyond those surveyed in the present study. Dissecting the mechanisms governing substrate selection and transport by SFXN1 and other mitochondrial carriers in specific physiological and pathological contexts, and how this is coordinated with the rest of the MC system, is necessary to ultimately decode the mitochondrion’s complex regulatory network.

## ACKNOWLEDGMENTS

We are grateful to Dr. Valeria Culotta and Dr. Sabrina Schatzman for suggestions regarding iron measurements. We thank Dr. Heather Neu of the Mass Spectrometry Center at the University of Maryland School of Pharmacy for processing and analyzing the samples for ICP-MS, Dr. Pete Pederson for the ATP5B antisera, and Dr. Adam Cornish for technical assistance. This work was supported in part by NIH grants (R01HL108882 to S.M.C.; R01NS072241 and R01DK116746 to M.J.W., and Biochemistry, Cellular, and Molecular Biology Program Training Grant T32GM007445 to Y.L.), a NSF grant (MCB-1330803 to C.F.C.), and pre-doctoral fellowships from the American Heart Association (16PRE31140006 to M.G.A. and 12PRE11910004 to Y.L.).

## AUTHOR CONTRIBUTIONS

M.G.A. and S.M.C. conceptualized the research, interpreted results, and wrote the manuscript, with input and approval from all authors; M.G.A. designed and conducted most of the experiments; E.S.S.A. designed, performed, and analyzed the metabolite quantification and isotope tracing experiments; S.R. performed the SILAC studies; L.F.R. performed and analyzed the CoQ measurements; Y-W.L. performed experiments for antibody generation; S.M.C., M.J.W., A.P. and C.F.C. contributed to the experimental design and data analysis and provided essential reagents.

## METHODS DETAILS

### Cell culture

Flp-In 293 cells (Thermo FIsher Scientific) were grown in DMEM with 4.5 g/L glucose (Corning), supplemented with 10% fetal bovine serum (Seradigm, Gibco), 2 mM L-glutamine (Gibco), and 100 µg/mL zeocin (Thermo FIsher Scientific), and kept at 37**°**C in 5% CO_2_. HeLa cells (ATCC) were maintained in the same media but without zeocin. For galactose-containing media, DMEM without glucose (Gibco) was supplemented with 10 mM galactose (Acros Organics), 10% FBS and 2 mM L-glutamine.

To make media lacking serine, DMEM without glucose, glutamine, serine, glycine, and sodium pyruvate (US Biological) was supplemented with 25 mM glucose (RPI), 4 mM glutamine, 30 mg/L glycine (RPI), sodium bicarbonate (Acros Organics), and 10% heat-inactivated dialyzed FBS (Gemini Bio, Thermo FIsher Scientific). For full media, 42 mg/L serine (RPI) was additionally included. 15 µM hemin (stock always freshly prepared in 20 mM NaOH) or 1 mM formate was supplemented as indicated.

### Cell proliferation

Cells were seeded in 6-well plates (2×10^5^ cells/well) in glucose-based media and allowed to adhere overnight. Per genotype, 6 wells were counted to monitor initial cell count at day of treatment (Day 1). Cells in another 6-well plate were washed twice with 1X PBS prior to switch to treatment media. Cells were counted at Day 5 by staining with trypan blue solution (Gibco) and using the Countess II FL automated cell counter (Thermo Fisher Scientific). Data were presented as cell count at Day 5 relative to Day 1 of treatment.

### Knock-out cell line generation via CRISPR/Cas9

Two guide RNAs (gRNAs) that target exon 2 of human SFXN1 were cloned separately into PX459, a gift from Feng Zhang (Addgene) (Ran et al., 2013). Cells were seeded in 6-well plates and grown in DMEM with 4.5 g/L glucose, 10% FBS and 2 mM L-glutamine, without antibiotics, and transfected with either construct using FuGENE HD Transfection Reagent (Promega) according to manufacturer’s protocol. HEK293 cells were transfected with gRNA #1 or #2 while HeLa cells were transfected with gRNA #2. After 24 hr, the medium was replaced with one containing the same components mentioned, but further supplemented with 1 mM sodium pyruvate (Gibco) and 50 µg/mL uridine (Sigma-Aldrich) (which were subsequently found to be not essential additions). 48 hr post-transfection, cells were selected with 2 µg/mL puromycin (Gibco) for 2 days, then grown in puromycin-free medium until they reached confluency. They were passaged and a portion was used for genomic DNA extraction using the Puregene Core Kit A (Qiagen) and T7 endonuclease assay (Genecopoeia) to test if indels were introduced. The remaining cells were used for seeding at low density in 10-cm dishes. Single colonies were isolated by ring cloning, expanded, and analyzed by immunoblotting with SFXN1 antibody to select KO clones.

gRNA that targets human AGK, under the U6 promoter, was synthesized as a gene block fragment (Integrated DNA technologies) and cloned by In-Fusion cloning (Clontech) into the pEF6-nls-YFP-2A-Cas9 vector, as previously detailed (Bowman et al., 2017). Blasticidin (7.5 µg/mL; InvivoGen) selection was done 48 hr after transfection and carried out for two weeks. YFP-positive clones were selected by flow cytometry and surviving clones were screened for the absence of AGK by genotyping and immunoblotting.

### Genotyping of SFXN1- and AGK-null clones

Genomic DNA was extracted from WT and KO clones. The region surrounding the gRNA target site was amplified by PCR, followed by digestion with HindIII/SacII (NEB), and ligation into pBSK(-). At least 10 individual transformants were analyzed by sequencing to determine the mutations introduced. Nucleotide sequence alignments were performed by Clustal Omega.

### Molecular Biology

Human SFXN1 cDNA was amplified from HEK293 mRNA by using GoScript™ Reverse Transcription Mix, Oligo(dT) (Promega). Human AGK was copied from HeLa cDNA. CNAP-SFXN1, SFXN1-HA, and AGK^G126E^ were generated by overlap-extension polymerase chain reaction (PCR) using Platinum™ *Pfx* DNA polymerase (Thermo Fisher Scientific). All were cloned into pcDNA5/FRT (Invitrogen) and sequenced before use for transfection. To purify recombinant AGK and recombinant ANT2, human AGK and ANT2 were cloned into pET28a vector (Novagen) containing a N-terminal hexahistidine tag. Protein induction, extraction and affinity purification were carried out as described previously (Lu et al., 2017).

### Whole cell extraction

Cells were washed with ice-cold 1X PBS and incubated in RIPA buffer (20 mM HEPES-KOH, pH 7.4, 1% (v/v) Triton X-100, 50 mM NaCl, 1 mM EDTA, 2.5 mM MgCl_2_, 0.1% (w/v) SDS) with 1 mM PMSF for 30 min at 4°C, with constant agitation. Soluble and insoluble materials were separated by centrifugation at 21,000*xg* for 30 min at 4°C. The supernatant was transferred to a fresh tube, and protein quantification was performed with Pierce™ BCA Protein Assay Kit (Thermo FIsher Scientific).

### Mitochondrial isolation

Cells were seeded onto 150 mm x 25 mm culture dishes and expanded. At around 80% confluency, glucose-based medium was changed to galactose-based medium and the cells were further grown for 48 hr. For supplementation experiments, the galactose-containing media was supplemented with 15 µM hemin, 1 mM formate, 10 mM dimethyl α-ketoglutarate (DMK; Sigma-Aldrich), 2 mM serine, 2 mM glycine, 2 mM alanine (RPI), or 2 mM cystine dihydrochloride (RPI), as indicated. For the minus (–) serine panel, cells were seeded into glucose-based medium and allowed to adhere for 48 hr before washing with 1X PBS and adding the indicated medium.

Mitochondrial isolation was performed based on a previous protocol (Frezza et al., 2007). Briefly, cells were washed with ice-cold 1X PBS and resuspended in 3 mL IBc buffer (200 mM sucrose, 10 mM Tris-MOPS, 1 mM EGTA/Tris, pH 7.4). Homogenization was performed using a Teflon Potter-Elvehjem motor-driven dounce set at 1600 rpm. Lysates were centrifuged twice at 600*xg* for 10 min at 4°C to remove cellular debris and the nuclear fraction. The supernatant was then spun down at 7,000*xg* for 10 min at 4°C. The resulting pellet was resuspended in IBc buffer and centrifuged again at 7,000*xg* for 10 min at 4°C. An additional centrifugation at 10,000*xg* for 10 min at 4°C was performed and the pellet, which represents the crude mitochondria, was resuspended in IBc buffer to a final concentration of 10mg/mL. If not used right away, mitochondrial extracts were aliquoted, snap frozen in liquid nitrogen, and stored at -80°C.

### Immunoblotting

Equal protein amounts (20-50 µg depending on samples or antibodies used) were loaded onto SDS-PAGE gels (12% or 10-16%) and immunoblotting was carried out as described previously (Lu et al., 2016). Images were captured using Fluorchem Q (Cell Biosciences, Inc.) quantitative digital imaging system. Antibodies used for this study were as follows: AGK (AB Clonal Technology; this study), ALAS1 (Abcam; ab154860), ANT1 (Lu et al., 2017), ANT2 (Pacific Immunology; this study), ANT2/3 (Panneels et al., 2003), ATP5B (Peter Pedersen, Johns Hopkins University), β-actin (Sigma-Aldrich; A5441), COX4 (Abcam), CPOX (Proteintech), CYTB (Proteintech), CYTC (BD Biosciences), DNAJC19 (Proteintech), FECH (Proteintech), GRP75 (Antibodies Inc.), HA (Sigma-Aldrich), NDUFB6 (Abcam), NDUFS1 (Abcam), OPA1 (BD Biosciences), Protein C (Genscript), PHB1 (Biolegend), PHB2 (BioLegend), SDHA (Invitrogen, Abcam), SDHB (Abcam), SFXN1 (Proteintech), TIM23 (BD Biosciences), TOM20 (Santa Cruz Biotechnology), UQCRC2 (Abcam), and UQCRFS1 (Abcam). Goat anti-rabbit (Pierce) or mouse (Pierce) secondary antibodies conjugated to horseradish peroxidase were utilized. Images were minimally processed in Adobe Photoshop and assembled in Adobe Illustrator. Densitometric band analysis for protein quantitation was performed using Quantity One 1-D Analysis Software (Bio-Rad).

### Membrane association assays

HEK293 WT mitochondria (250 µg) were resuspended in 1 mL cold 5 mM Tris pH 7.5, incubated on ice for 30 min, and centrifuged at 21,000*xg* for 10 min at 4°C. The pellet was resuspended in 0.5 mL Sonication Buffer (0.6 M sucrose, 3 mM MgCl_2_, 20 mM HEPES-KOH, pH 7.4), sonicated thrice for 10 sec, with 30 sec intervals on ice, and spun down at 21,000*xg* for 10 min at 4°C. Ultracentrifugation (Optima MAX, TLA-120.1 rotor) at 175,000*xg* for 30 min at 4°C was carried out for the supernatant. The resulting soluble fraction was subjected to precipitation using 20% (v/v) TCA and 0.07% (w/v) sodium deoxycholate, while the pellet was directly resuspended in 1:1 2X RSB: 0.1 N NaOH. Analysis was done by SDS-PAGE and immunoblotting.

For carbonate extraction, 100 µg HEK293 mitochondria was resuspended in 0.5 mL of 0.1 M Na_2_CO_3_ of varying pH (pH 10.5, 11.0, 11.5, 12.0, 12.5), vortexed and incubated on ice for 30 min. Samples were subjected to ultracentrifugation and downstream steps as detailed above.

### Digitonin-based mitochondrial fractionation

The protocol used was adapted from a previous study (Lu et al., 2016). 100 µg mitochondria were resuspended in 50 µL SEHK buffer (250 mM sucrose, 5 mM EDTA, pH 7.0, 10 mM HEPES-KOH, pH 7.4, 200 mM KCl) supplemented with 0, 0.1, 0.2, 0.3, 0.4 or 0.5% (w/v) digitonin and 1 mM PMSF. Samples were briefly vortexed on low and incubated on ice for 2 min. 450 µL ice-cold SEHK buffer was then added to halt solubilization. Samples were fractionated by ultracentrifugation at 100,000*xg* for 10 min at 4°C. The resulting supernatants were subjected to TCA precipitation. Pellets from membrane and soluble fractions were resuspended in equal volumes of reducing sample buffer and analyzed by SDS-PAGE.

### Protease protection assay

To determine the submitochondrial localization of endogenous SFXN1, CNAP-SFXN1 (CNAP consists of a protein C and His_10_ tags) or SFXN1-HA, 200 µg intact mitochondria, mitoplasts, or solubilized mitochondria were treated with 125 µg/mL Pronase E (Sigma-Aldrich) for 30 min on ice, as indicated. Mitoplasting and solubilization was done by osmotic swelling in 5 mM Tris-HCl pH 7.5 or by utilizing 0.5% (v/v) sodium deoxycholate, respectively. Protease digestion was halted by adding 5 mM PMSF. Samples were TCA-precipitated and heated at 65°C for 5 min. This was followed by incubation on ice for 1 hr, and spinning at 21,000*xg* for 10 min at 4°C. Pellets were washed with 1 mL cold acetone, dried, resuspended in 1:1 2X RSB: 0.1 N NaOH, and boiled for 5 min. Analysis was done by SDS-PAGE and immunoblotting.

### Blue native (BN)-PAGE analysis

Mitochondria (100 or 150 µg for 1D, 150 µg for 2D, 150 µg for Coomassie staining) were solubilized in lysis buffer (20 mM Tris-Cl, 10% (v/v) glycerol, 100 mM NaCl, 20 mM imidazole, 1 mM CaCl_2_, pH 7.4) containing either 1% (w/v) digitonin (Biosynth International) or 1% (w/v) n-dodecyl-β-maltoside (DDM) (Thermo Fisher Scientific), and supplemented with protease inhibitors (1 mM PMSF, 2 µM pepstatin A, 10 µM leupeptin). Isolates were clarified by spinning at 21,000*xg* for 30 min at 4°C. Supernatants with BN sample buffer added were resolved using in-house-made 4-16% BN-PAGE gels for immunoblotting, or 6-16% BN-PAGE gels for Coomassie staining (0.25% (w/v) Coomassie Brilliant Blue R250, 40% methanol, 10% glacial acetic acid). For 1D BN-PAGE, the steps following transfer were adapted as previously described (Maio et al., 2017). Briefly, the PVDF membrane was soaked in 8% acetic acid for 20 min, rinsed with water, and air-dried. The membrane was washed with methanol until most of the residual Coomassie Blue G-250 was removed, before proceeding to standard immunoblotting procedures. The 2D BN/SDS-PAGE analysis was performed as previously described (Claypool et al., 2008).

### SILAC

HEK293 WT and two *AGK* KOs were subjected to three-plex SILAC. For set 1, WT was grown in “light” media, *AGK* KO1 in “medium” media, and *AGK* KO2 in “heavy” media. For set 2, the media for *AGK* KO1 and KO2 were swapped. DMEM without lysine, arginine, and sodium pyruvate (Gibco) was the base for all three media types. Light media was prepared by supplementing the base media with unlabeled L-lysine:2HCl (0.8 mM) (Acros Organics) and L-arginine:HCl (0.4 mM) (Acros Organics). To the same final concentrations, medium and heavy media contained added ^2^H_4_-L-Lysine:2HCl and ^13^C_6-_L-Arginine:HCl, or ^13^C_6_^15^N_2_-L-Lysine:2HCl and ^13^C_6_^15^N_4_-L-Arginine:HCl (Cambridge Isotope Laboratories), accordingly. Cells were passaged in their respective medium at least five times before initiating the experiment to ensure that the isotopic amino acids were incorporated. Two days before mitochondrial isolation, cells were cultivated in medium with the same formulation except that glucose was replaced by 10 mM galactose. A preliminary MS run was carried out to make sure that isotope labeling was >95% before proceeding to the actual experiment. Equal concentrations of mitochondria as determined by BCA assay were mixed and processed for liquid chromatography-tandem mass spectrometry (LC-MS/MS), as previously described (Wu et al., 2014). For each sample, 1 µg was injected twice in an Orbitrap Fusion Lumos mass spectrometer coupled to an Easy nanoLC 1200 ultra-high pressure liquid chromatography (UPLC) system (Thermo Scientific). Peptides were separated on an analytical column (50 cm x 75 µm, PepMap C_18_ 1.9 µm, 100 Å, EasySpray, Thermo Scientific) using a linear gradient of 5-30% solvent B (acetonitrile in 0.1% (v/v) formic acid) over 85 min with total run time of 120 min. Both MS and MS/MS were measured using an Orbitrap mass analyzer in top speed mode. Raw data was searched using Sequest algorithm against Human protein database (NCBI RefSeq 85) in Proteome Discoverer Software Suite v2.0 (Thermo Scientific). The search parameters included a maximum of two missed cleavages; carbamidomethylation at cysteine as a fixed modification; N-terminal acetylation, oxidation at methionine, SILAC labeling ^13^C_6_,^15^N_2_-lysine, ^2^H_4_-lysine, ^13^C_6_-arginine and ^13^C_6_,^15^N_2_-arginine as variable modifications. Precursor tolerance was set to 10 ppm and MS/MS tolerance to ±0.02 Da. False discovery rate was set to 1% at the peptide-spectrum matches (PSMs), peptide and protein level. Data have been deposited to the ProteomeXchange Consortium via the PRIDE partner repository with the dataset identifier PXD019826.

### Iron chelation

Cells were depleted of iron by treatment with deferoxamine mesylate (DFO) (Sigma-Aldrich), an iron chelator. Cell viability of HEK293 WT and *SFXN1* KOs in different DFO concentrations was assessed using CellTiter-Glo® 2.0 (Promega). 1×10^4^ cells were seeded per well in a white-walled microplate and allowed to adhere overnight before DFO treatment for 24 hr. Data were presented as luminescence values relative to no treatment.

### Metal level quantification

Whole cell or mitochondrial pellets were snap-frozen until analysis by inductively-coupled mass spectroscopy (ICP-MS). Samples were warmed to room temperature and centrifuged for 2 min at 14,000*xg*. To each sample, 200 µL of concentrated HNO_3_ was added, the samples vortexed and then incubated at 80°C overnight. Samples were vortexed for 30 sec while still warm, and diluted with 2 mL of milliQ water, transferred to 15 mL conical tubes, and analyzed by ICP-MS (University of Maryland School of Pharmacy Mass Spectrometry Center).

### Enzyme activity measurements

Activities of Complex I (NADH dehydrogenase), Complex II (succinate dehydrogenase), and Complex IV (cytochrome c oxidase) were determined using Abcam’s Complex I (Abcam), Complex II (Abcam), or Complex IV (Abcam) Human Enzyme Activity Microplate Assay Kits, accordingly. Mitochondria (50 µg for Complex I and II, 10 µg for Complex IV) were solubilized at 5 mg/mL in the provided detergents supplemented with 1 mM PMSF based on manufacturer’s recommendations. Complex I activity was measured by monitoring the increase in absorbance at 450 nm which is coupled to the oxidation of NADH to NAD^+^. Complex II activity was determined as the rate of decrease in absorbance at 600 nm, when ubiquinol is generated and causes the reduction of DCPIP. Complex IV activity was quantified based on cytochrome c oxidation, which leads to a decrease in absorbance at 550 nm.

Complex III activity was measured as previously documented (Tzagoloff et al., 1975). Briefly, mitochondria were solubilized in reaction buffer (50 mM KP_i_, 2 mM EDTA, pH 7.4) containing 0.5% (w/v) DDM, or 0.5, 1.0, or 1.5% (w/v) digitonin, and spiked with protease inhibitors. 30 µg was added to reaction buffer with 0.08% (w/v) equine heart cytochrome c (Sigma-Aldrich) and 1 mM potassium cyanide. The reaction was started by adding 100 µM decylubiquinol. The reduction of cytochrome c was recorded as the rate of absorbance increase at 550 nm.

Alanine aminotransferase (ALT) activity was analyzed using the ALT Assay Kit (Sigma-Aldrich). 1×10^6^ cells were homogenized in 200 µL ice-cold ALT assay buffer. Samples were spun at 15,000*xg* for 10 min at 4°C. Per reaction, 20 µL of the supernatant was used. The initial fluorescent intensity was measured (λ_ex_/λ_em_= 535/587 nm), then the samples were incubated at 37**°**C with a reading taken every 5 min for 1 hr. Calculations were performed per manufacturer’s protocol. Enzyme activities were normalized to protein concentrations of remaining supernatants as determined by BCA assay.

Glutamate dehydrogenase (GDH) activity was analyzed using the GDH Assay Kit (Sigma-Aldrich). 1×10^6^ cells were lysed in 200 µL ice-cold GDH assay buffer. Samples were incubated on ice for 10 min prior to centrifugation at 13,000*xg* for 10 min at 4**°**C. Per reaction, 50 µL of the supernatant was used. Samples were incubated at 37**°**C for 3 min and initial absorbance was measured at 450 nm. Further incubation at 37**°**C took place for 1 hr, with readings every 5 min. Calculations were performed per manufacturer’s protocol. Enzyme activities were normalized to protein concentrations of remaining supernatants as determined by BCA assay.

### Gene expression analysis by quantitative PCR (qPCR)

Total RNA extracts were prepared using PureLink® RNA Mini Kit and DNAse set (Invitrogen) according to manufacturer’s protocol. 25 ng RNA was used as template in 20 µL reactions for qPCR using EXPRESS One-Step SYBR GreenER kit (Thermo FIsher Scientific), with ROX as reference dye, and 200 nM each of forward and reverse primers for genes of interest. No template and no reverse transcriptase controls were included and samples were analyzed in duplicate. Reactions were run using the QuantStudio 6 Flex Real-Time PCR System (Applied Biosciences) with the following conditions: (a) 5 min at 50°C and 2 min at 95°C (cDNA synthesis), (b) 15 sec at 95°C and 1 min at 60°C (40-cycle amplification), and (c) melt curve analysis. Gene expression was expressed as d*C*_*T*_ or fold-change (FC) relative to WT, calculated based on the 2^−ΔΔ^*C*_*T*_ method, with GAPDH as the reference gene.

### mtDNA content

Genomic DNA was isolated using Gentra Puregene Cell Kit (Qiagen). qPCR was carried out with FastStart Universal SYBR Green Master Rox (Roche) according to manufacturer’s guidelines, with 10 ng gDNA as template in a 20-µl reaction mixture. QuantStudio 6 Flex Real-Time PCR System was used. *ND1* is a mtDNA-encoded subunit of Complex I whereas β2M serves as a nuclear-encoded reference gene.

### Translation of mitochondrial-encoded proteins

*In vivo* labeling of mitochondrial translation products was carried out based on a previous protocol (Sasarman and Shoubridge, 2012). Cells seeded on 6-well plates were washed with warm 1X PBS, switched to labeling media (DMEM without methionine and cysteine (Gibco), 10% dialyzed FBS (Gibco), 1X GlutaMAX™ (Thermo Fisher Scientific), 110 mg/L sodium pyruvate (Gibco), and 2 mM L-Glutamine (Gibco)) and incubated at 37°C, 5% CO_2_ for 30 min. After washing with 1X PBS, cells were treated with 100 µg/mL anisomycin (Sigma-Aldrich) and incubated at 37°C, 5% CO_2_ for 5 min to inhibit mitochondrial translation. Per well, 240 µCi of EasyTag™ EXPRESS^35^S protein labeling mix (Perkin Elmer) was added. Labeling took place at 37°C, 5% CO_2_ for 30 min, after which the media was switched to media A (DMEM with 10% FBS, 2 mM L-glutamine and 1 mM sodium pyruvate) and the plates returned to the incubator for 10 min. Cells were then washed with 1X PBS thrice before collection, resuspension in 1X PBS with 1 mM PMSF, and protein quantification via BCA assay. Per sample, 50 µg were resuspended in gel loading buffer (93 mM Tris-Cl, pH 6.8, 7.5% (v/v) glycerol, 1% (w/v) SDS, 0.25 mg/mL bromophenol blue, 3% β-mercaptoethanol) and resolved on a 15-20% SDS-PAGE gel. Gel was stained with Coomassie solution, destained, vacuum-dried, and exposed to Imaging K-screen (Bio-Rad). Radiolabeled proteins were visualized by autoradiography (Pharos FX, Quantity One 1-D Analysis Software).

### Oxygen consumption rate (OCR) and extracellular acidification rate (ECAR) measurements

The XF^e^96 extracellular flux analyzer (Agilent) was utilized to record OCR and ECAR. For analyzing mitochondrial respiration and glycolytic parameters in intact cells, the XF Cell Mito Stress Test (Agilent) and Glycolysis Stress Test (Agilent) kits were used. Cells were seeded at 10,000 cells/well in a Seahorse XF96 V3 PS cell culture microplate (Agilent) coated with 0.001% (w/v) poly-L-lysine (Sigma-Aldrich) for improved cell adherence. After 48 hr, cells were washed twice with XF base medium (Agilent) supplemented with 1 mM sodium pyruvate, 2 mM L-glutamine, and for the Mito Stress Test only, 10 mM glucose. Cells were then incubated in a humidified non-CO_2_ incubator at 37°C for a 1-hr degassing. For the Mito Stress Test, OCR values were recorded under basal conditions, and after the addition of 2 µM oligomycin, 250 nM FCCP, or 1 mM rotenone/antimycin A. For the Glycolysis Stress Test, ECAR values were obtained under basal conditions, and after the addition of 10 mM glucose, 2 µM oligomycin or 50 mM 2-deoxyglucose.

To measure the dependence of cells on the oxidation of glucose/pyruvate, glutamine or fatty acids, inhibitors of pathways utilizing these substrates were used: 2 µM UK-5099 (Tocris), 3 µM BPTES (Tocris) and 4 µM etomoxir (Tocris) inhibit the mitochondrial pyruvate carrier (MPC), carnitine palmitoyltransferase 1A (CPT1A), and glutaminase (GLS), respectively. Cells were seeded at 12,500 cells/well and allowed to adhere for 48 hr. After media exchange with XF base medium supplemented with 2 mM L-glutamine and 10 mM glucose, without sodium pyruvate, cells were incubated in a humidified non-CO_2_ incubator at 37°C for 1 hr. Baseline OCR was recorded after 24 min, then the inhibitor of the targeted pathway was injected. Target inhibitor OCR was recorded after 48 min, and then the inhibitors of the two remaining pathways were injected. OCR was monitored following an additional 48 min incubation in presence of all three inhibitors. % dependency is calculated as: (baseline OCR-target inhibitor OCR)/(baseline OCR-all inhibitors OCR).

After each experiment, media was removed and the plate was frozen at -20°C in preparation for DNA content quantification with CyQUANT Assay (Thermo Fisher Scientific). Calculations were performed per manufacturer’s instructions, and OCR and ECAR measurements were normalized to DNA content.

For OCR measurements in isolated mitochondria, fresh extracts were diluted to 1 mg/mL in IBc buffer. For the electron flow assay (Figure 3E), 5 µg mitochondria were dispensed per well. To enhance mitochondrial adherence, the microplate was spun at 2,000*xg* for 20 min at 4°C. 50 µL of 1X MAS, pH 7.2 (220 mM D-mannitol, 70 mM sucrose, 10 mM KH_2_PO_4_, 5 mM MgCl_2_·6H_2_O, 2 mM K^+^ HEPES, pH 7.4, 1 mM EGTA, 0.2% (w/v) fatty acid-free BSA) with 10 mM pyruvate, 2 mM malate and 4 µM CCCP was added carefully to each well. Compounds reconstituted in 1X MAS without BSA were delivered to cartridge ports so as to have the following final concentrations after sequential injections: port A, 4 µM rotenone; port B, 10 mM succinate; port C, 4 µM antimycin A; port D, 10 mM ascorbate/100 µM TMPD. For the substrate utilization experiment (Figure 5C), the following were prepared in 1X MAS, pH 7.2, with 0.2% (w/v) fatty-acid free BSA: 10 mM pyruvate/2 mM malate, 10 mM glutamate/10 mM malate, 10 mM DMK/2 mM malate, 10 mM glutamine/2 mM malate, 5 mM succinate/10 µM rotenone, 10 mM glycerol-3-phosphate/2 µM rotenone, and 80 µM palmitoylcarnitine/2 mM malate. 5 µg mitochondria/well was used except when palmitoylcarnitine/malate served as substrate, wherein 10 µg was utilized instead. After allowing the mitochondria to adhere to the plate by centrifugation, 50 µL of 1X MAS with BSA containing the substrates of interest was added per well. Injections were as follows, with the final concentrations indicated: port A, 4 mM ADP; port B, 2 µM oligomycin; port C, 4 mM CCCP; 4 µM rotenone/antimycin A.

### NAD^+^/NADH and NADPH/NADP^+^ measurements

Cellular NAD^+^/NADH and NADPH/NADP^+^ were determined using the NAD/NADH-Glo™ Assay (Promega) and NADP/NADPH-Glo™ Assay (Promega), accordingly. 5 × 10^5^ cells were seeded into 6-well plates and incubated at 37°C, 5% CO_2_, for 2 days. On the day of the experiment, cells were washed with 1X PBS and 400 µL of ice-cold lysis buffer (1:1 1X PBS:1% (w/v) dodecyltrimethylammonium bromide (DTAB; Sigma-Aldrich) in 0.2 N NaOH) added. Lysates were frozen at -80°C prior to the assay, which was performed as previously described (Sullivan et al., 2015).

### CoQ measurements

HEK293 cellular CoQ content was measured as previously described (Fernandez-Del-Rio et al., 2017). In brief, cell pellets were resuspended in 1X PBS, a portion of which was used for protein quantification by BCA assay while the remaining was subjected to lipid extraction using methanol and petroleum ether containing the internal standard dipropoxy-CoQ_10_. CoQ_9_ and CoQ_10_ standards containing dipropoxy-CoQ_10_ were also prepared and lipid extracted to generate standard curves. RP-HPLC MS/MS was employed to measure CoQ_9_ and CoQ_10_ levels, with the reconstituted lipid extracts first separated on a Luna 5 µM PFP(2) 100A column (100 x 4.6 mm, 5 µm; Phenomenex) in 90% solvent A (95:5 methanol/isopropanol, 2.5 mM ammonium formate) and 10% solvent B (isopropanol, 2.5 mM ammonium formate) at a constant flow rate of 1 mL/min. Quantifications were based on total peak areas normalized against the internal standard dipropoxy-CoQ_10_, calculated using the CoQ_9_ and CoQ_10_ standard curves, and normalized to protein content.

### Metabolite quantifications

TCA metabolite measurements were obtained via LC-MS/MS. HEK293 WT and *SFXN1* KOs were seeded at a density of 2×10^5^ cells/well in 24-well plates. After 48 hr, cells were washed twice with ice-cold 1X PBS, and extracted with ice-cold 80:20% (v/v) methanol:MS-grade water. Samples were kept on dry ice during the collection process. Samples were then vortexed vigorously and centrifuged at 12,000 rpm for 15 min at 4°C. Pellets were used for protein quantification while supernatants were dried overnight by vacuum centrifugation.

At the time of analysis, dried samples were reconstituted with water containing 0.2% (v/v) formic acid. TCA cycle intermediates were separated on Nexera UHPLC (Shimadzu) using X-Terra C18 UHPLC column (2.6um, 50mm, 2.1mm; Waters, Milford, MA) using a previously published method with a minor change in gradient (Al Kadhi et al., 2017). 0.2% (v/v) formic acid in water and 0.2% (v/v) formic acid in acetonitrile were used as mobile phases A and B, respectively. Total flow was set to 0.2 mL/min, with a linear gradient applied as follows: starting condition 0% B was linearly increased to 5% B in 4 min; 2 min at 100% B was added for column washing purposes. In between samples, the column was re-equilibrated for 3 min. Sample injection volume was 2 μL. Samples were kept at 4°C, and oven temperature was set to 21°C. Eluates were detected using AB SCIEX 4500 triple quadrupole instrument. Mass spectroscopy parameters were set for each metabolite by direct infusion of corresponding standards. Instrument was operated under following conditions: Curtain gas, 20V; Temperature, 500°C; Ion source gas 1, 50V; Ion source gas 2, 40V; Ion spray voltage, -4500V; CAD gas, Low; Entrance Potential, -10V. MultiQuant (AB SCIEX) was used to profile peaks and peak area was used for metabolite profiling. Mixture of TCA standards was utilized for standard curve generation in the μM range.

For the glutamine or glucose labeling experiments, cells were incubated with 4 mM [U-^13^C]-glutamine (Cambridge Isotope Laboratories) or 25 mM [U-^13^C]-glucose (Cambridge Isotope Laboratories) for 3 hr prior to processing as described above. Isotopomers were quantified using MultiQuant and presented as % of total metabolite signal. Metabolite abundances were normalized to protein content.

Cellular glycine and serine concentrations were measured using the Glycine and Serine Fluorimetric Assay Kits (Biovision), accordingly, following the manufacturer’s protocol. Briefly, 1×10^6^ cells were used to generate lysates which were deproteinized using 10 kDa spin columns (Millipore Sigma) and 5 or 10 μL of the filtrate was used for the assay. Readings were normalized to protein concentration as determined by BCA assay.

### Heme content

Heme content was measured based of PPIX fluorescence as previously documented (Sassa, 1976; Sinclair et al., 2001), with some modifications. 1×10^5^ cells was resuspended in 500 µL 20 mM oxalic acid and kept in the dark at 4°C. After 24 hr of acid extraction, 500 µL of 2 M oxalic acid (Sigma-Aldrich) was added and the samples were mixed by pipetting. Half was transferred to new amber tubes, which were then incubated at 98°C for 30 min in order to remove iron from heme and liberate PPIX. The remaining half was kept at RT for the same duration. Samples were centrifuged at RT at 12,000*xg* for 2 min before transferring 200 uL of the heated or unheated samples in duplicate into black-walled 96-well plates. Fluorescence was measured using a CLARIOstar microplate reader (BMG Labtech), at excitation and emission wavelengths of 400 nm and 620 nm, respectively.

### Treatment with heme biosynthesis inhibitors

One day post-seeding in glucose-based media, cells were treated with 0, 125, 250 or 500 µM succinylacetone (SA; Sigma-Aldrich) or 10 mM N-methyl protoporphyrin IX (NMP; Santa Cruz) for 48 hr prior to whole cell extraction and immunoblotting for proteins of interest.

### Statistical analysis

Results are presented as means, with error bars representing the standard error of the mean. Unpaired student’s t-test was used to evaluate statistical significance of differences between means (\**p<0.05*, \*\**p<0.01*, \*\*\**p<0.001*).

### Oligonucleotides used in this study

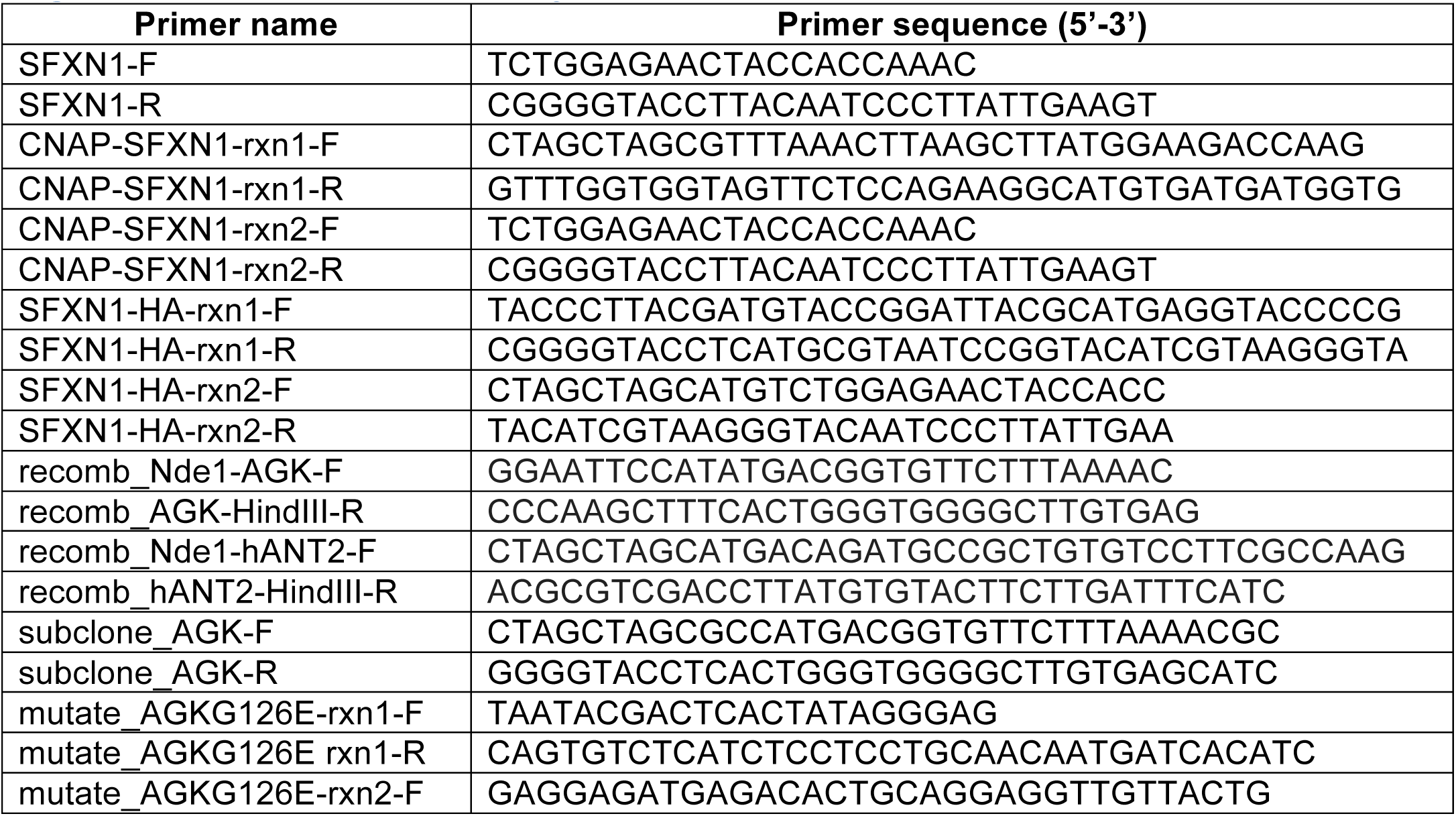
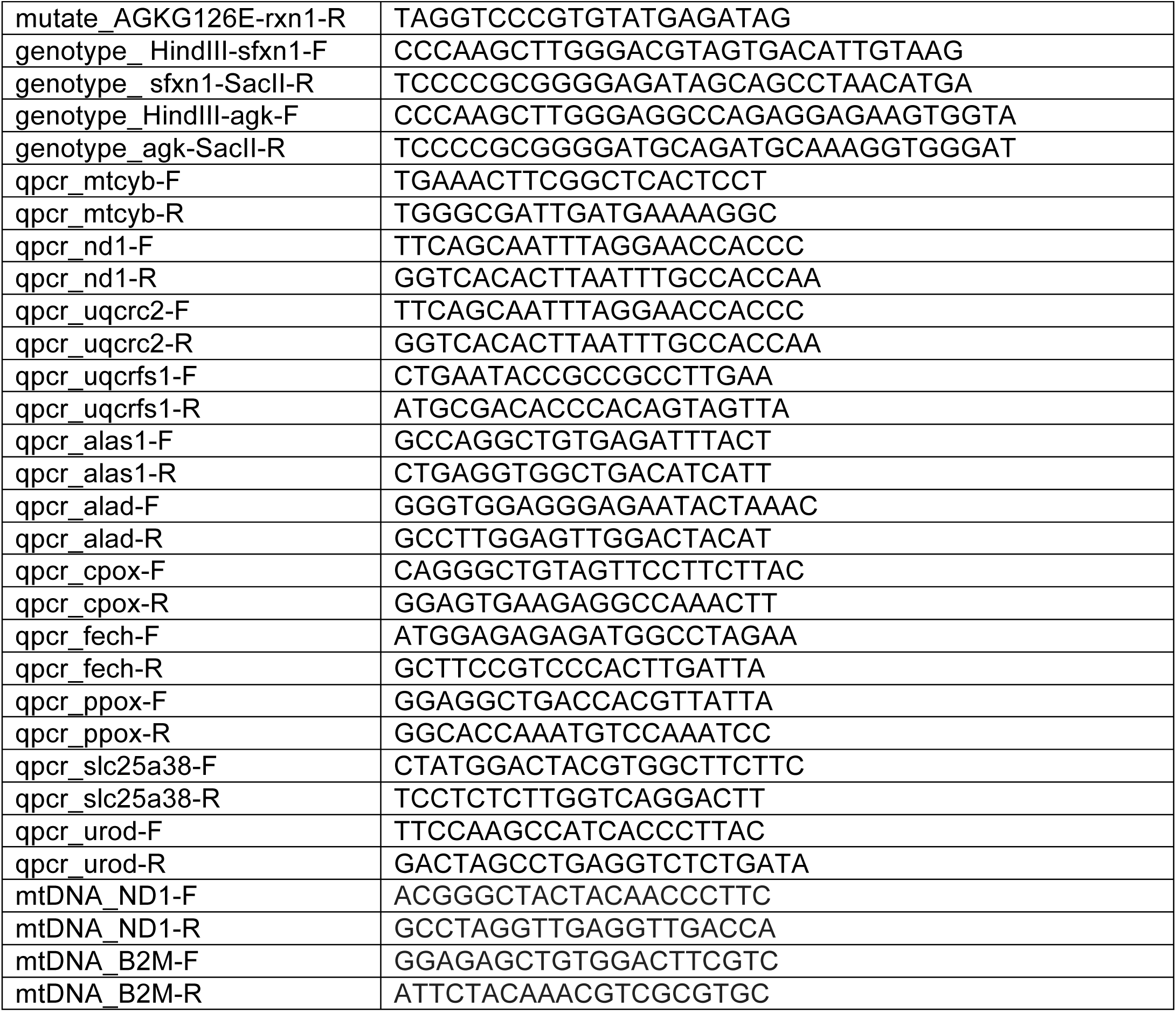

---

**Figure S1 related to Figure 1.**
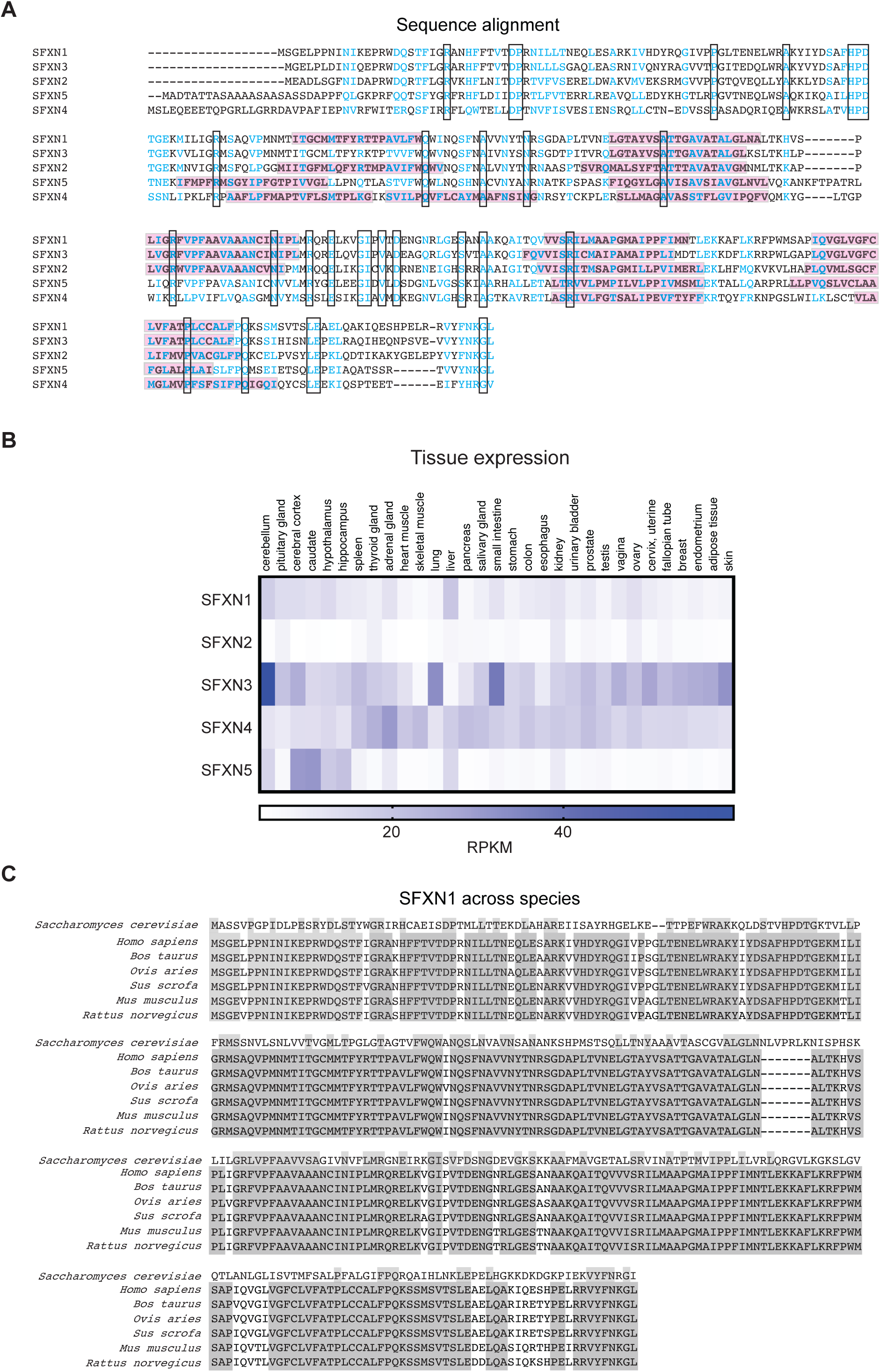
SFXN isoforms. (A) Alignment of SFXN isoforms. Identical amino acids are boxed whereas highly similar ones are highlighted blue. Predicted transmembrane domains are in pink. (B) Tissue distribution of SFXN isoforms. Gene expression is reported as median RPKM, generated by the GTEx project and accessed via the Human Protein Atlas. (C) SFXN1 is highly conserved in eukaryotes. *Saccharomyces cerevisiae* possesses a related protein (Fsf1p).

**Figure S2 related to Figure 1.**
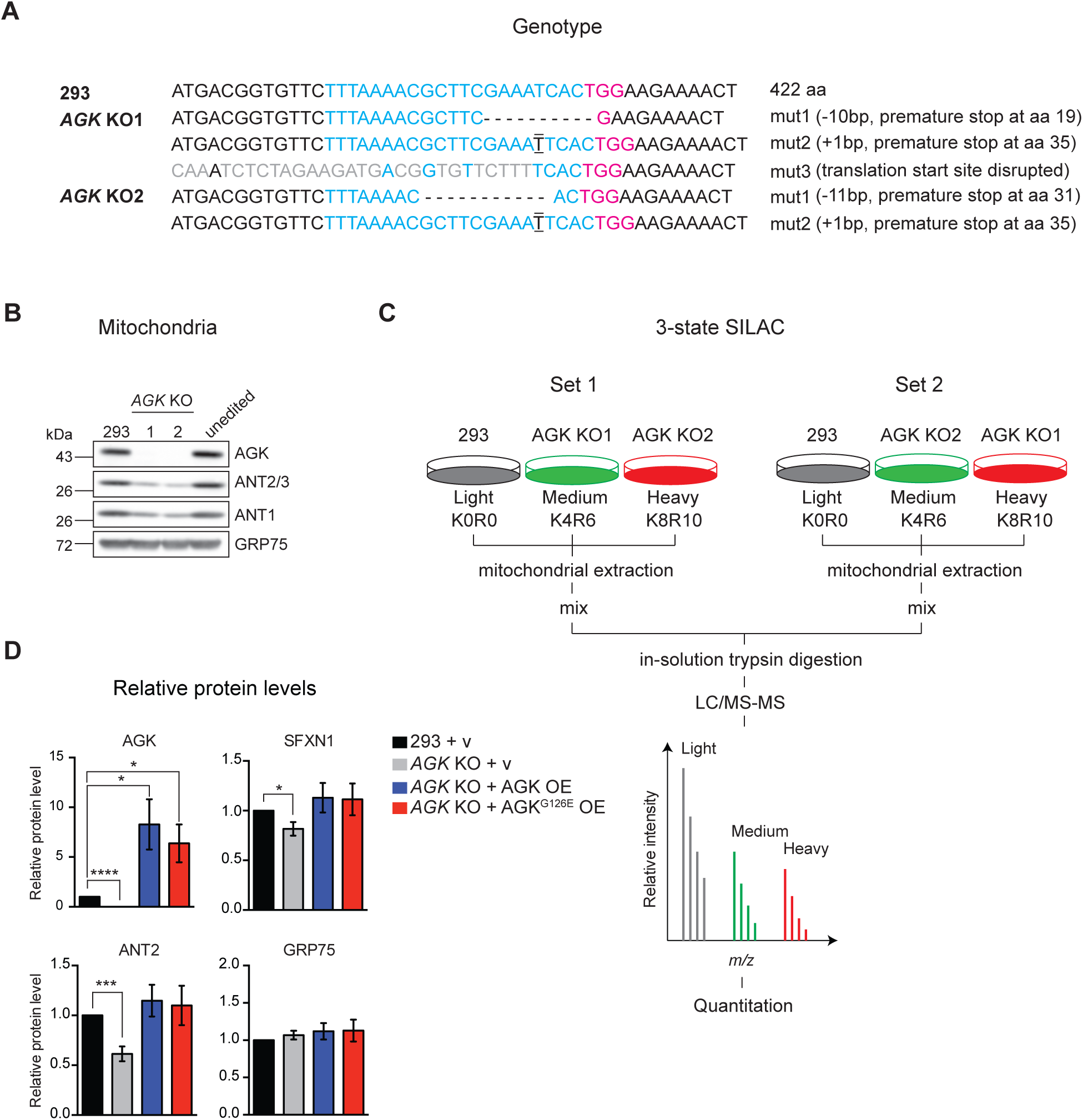
*AGK* disruption reduces steady-state protein levels of mitochondrial carriers including SFXNs. (A) Genomic mutations documented in two HEK293-based *AGK* KOs generated by CRISPR/Cas9. gRNA target site is in blue, while the protospacer-adjacent motif (PAM) sequence is in red. (B) Immunoblotting using mitochondrial isolates for AGK, ANT1, and ANT2/3 in WT, AGK-deficient clones, and an unedited clone that went through the transfection process with the CRISPR construct. GRP75 served as a loading control. (C) SILAC scheme related to Figure 1J. (D) Densitometric analysis of bands for proteins in Figure 1K. Protein levels in WT were set to 1.0. (mean±SEM, *n≥4*). \**p<0.05*, \*\**p<0.01*, \*\*\**p<0.001*, unpaired Student’s t-test

**Figure S3 related to Figures 2 and 3.**
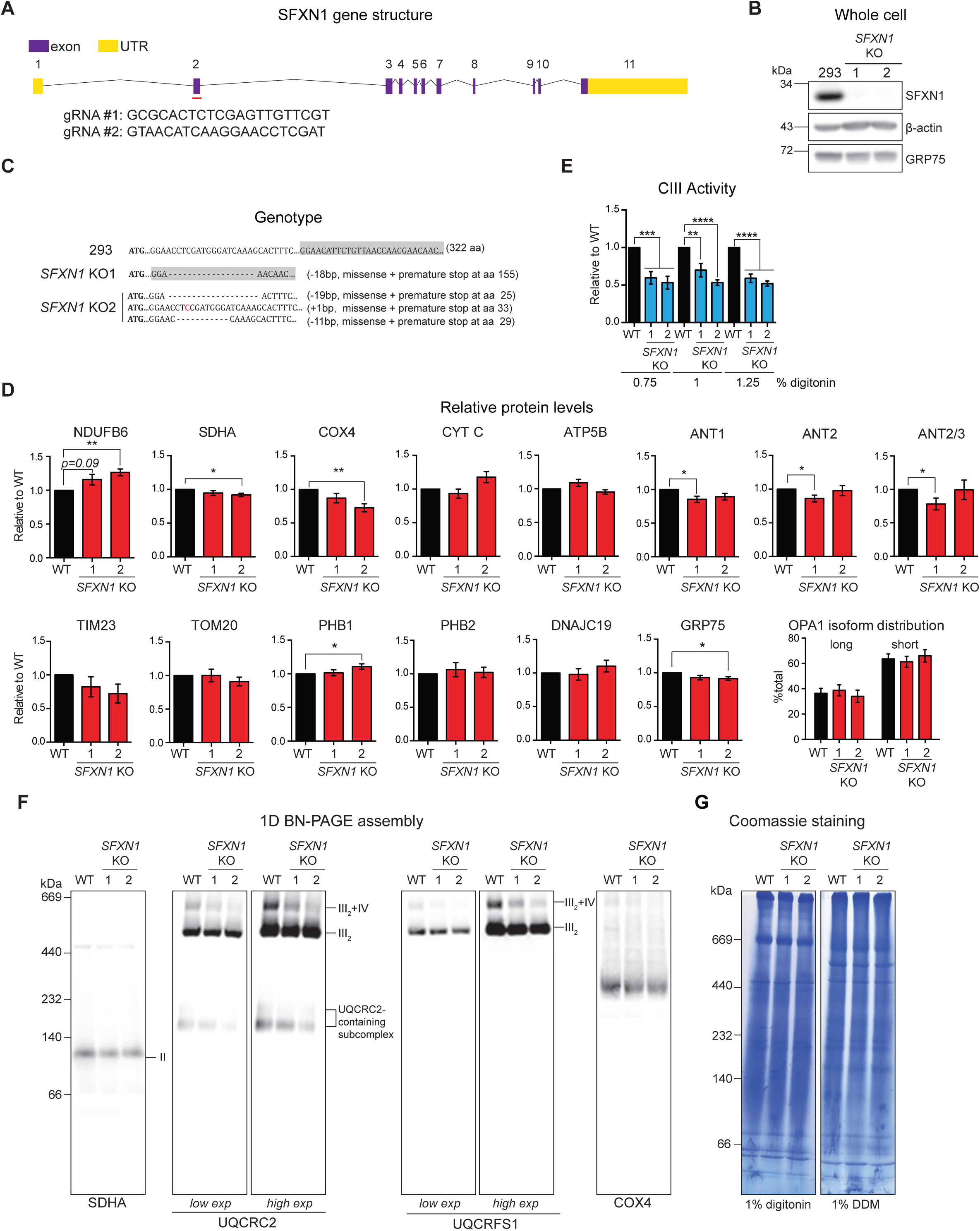
Effects of SFXN1 ablation on the levels, function and assembly of respiratory complexes. (A) *SFXN1* gene structure showing the target sites of gRNAs used for CRISPR/Cas9. (B) Whole cell extracts from WT and *SFXN1* KO clones were analyzed by SDS-PAGE and immunoblotting. (C) Genotyping shows genetic lesions resulting in premature stop codons in both clones. (D) Densitometric analysis of bands for other proteins immunoblotted as shown in Figure 2A. Protein levels in WT were set to 1.0. (mean±SEM, *n*≥4). (E) Complex III activity was measured by solubilizing mitochondria with increasing concentrations of digitonin and monitoring cytochrome *c* reduction at 550 nm (mean±SEM, *n*=6). (F) 1D BN assembly of respiratory complexes I-IV was assessed by solubilizing mitochondria in 1% (w/v) DDM, resolving in a 4-16% BN gel, and immunoblotting for the indicated complex subunits. (G) Coomassie staining of mitochondria solubilized with 1% (w/v) digitonin or 1% (w/v) DDM and resolved on 6-16% BN gel. \**p<0.05*, \*\**p<0.01*, \*\*\**p<0.001*, unpaired Student’s t-test

**Figure S4 related to Figure 3.**
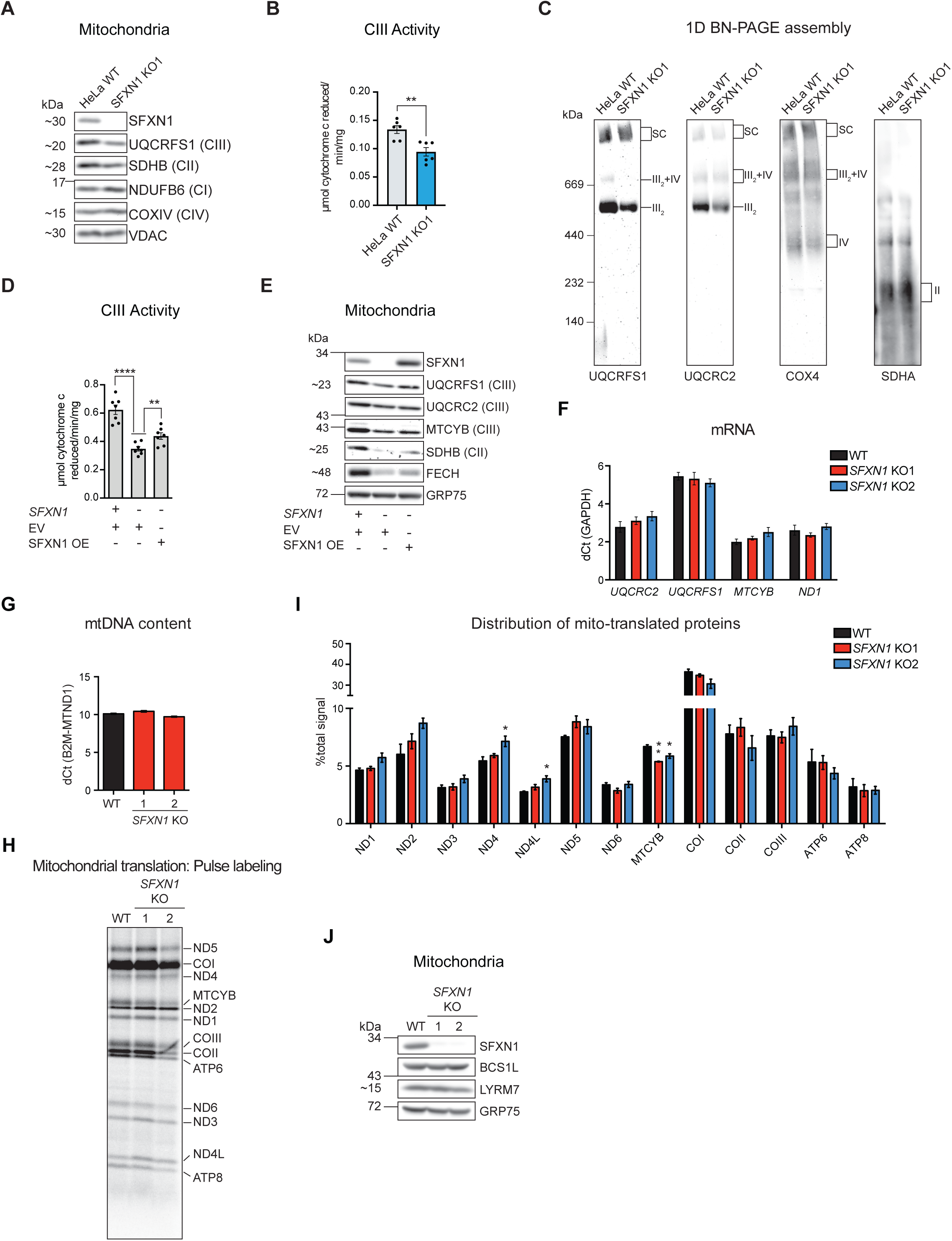
Complex III-associated function of SFXN1 does not derive from changes in mtDNA content, its transcription or translation, or altered CIII assembly factor levels. HeLa was used for (A) to (C). HEK293 was used for the remainder of the experiments. (A) Immunoblotting for select OXPHOS subunits using mitochondrial extracts. (B) Complex III activity was measured after solubilizing mitochondria with 0.5% (w/v) DDM and monitoring cytochrome *c* reduction at 550 nm (mean±SEM, *n*=6). (C) 1D BN assembly of Complexes II-IV was analyzed by solubilizing mitochondria in 1% (w/v) digitonin, resolving clarified extracts on a 4-16% BN gel, and immunoblotting for the indicated subunits. (D) Complex III activity was measured after solubilizing mitochondria with 0.5% (w/v) DDM and monitoring cytochrome *c* reduction at 550 nm (mean±SEM, *n*=7). (E) Immunoblotting for select mitochondrial proteins using mitochondrial isolates. (F) Gene expression analysis of CIII subunits *UQCRC2, UQCRFS1* and *MTCYB* by quantitative PCR (qPCR). *ND1*, a Complex I subunit, served as a control. Data presented as mean fold-change (FC)±SEM (*n*=6) relative to HEK293 WT. FC was calculated using the 2^*-ΔΔCt*^ method. GAPDH served as the housekeeping gene. (G) mtDNA content determination by qPCR. Genomic DNA isolated from the indicated cell lines served as template. mtDNA levels are presented as mean dCt±SEM (*n*=6) between the mitochondria-encoded *ND1* and the nuclear-encoded reference gene β-2-microglobulin (β2M). (H) Mitochondrial translation experiments. Cytosolic translation was inhibited by anisomycin (100 µg/mL), and mitochondrially-encoded polypeptides were pulse-labeled for 1 hr with ^35^S-methionine/cysteine (200 µCi/mL). Bands were detected by phosphoimaging. (I) Densitometric band analysis was performed and the distribution of signals from mitochondrially-translated products was calculated as a percentage of the total signal per experiment (mean % total signal±SEM, *n*=3). (J) Immunoblotting for CIII assembly factors using mitochondrial extracts. \**p<0.05*, \*\**p<0.01*, \*\*\**p<0.001*, unpaired Student’s t-test

**Figure S5 related to Figures 4, 6 and 7.**
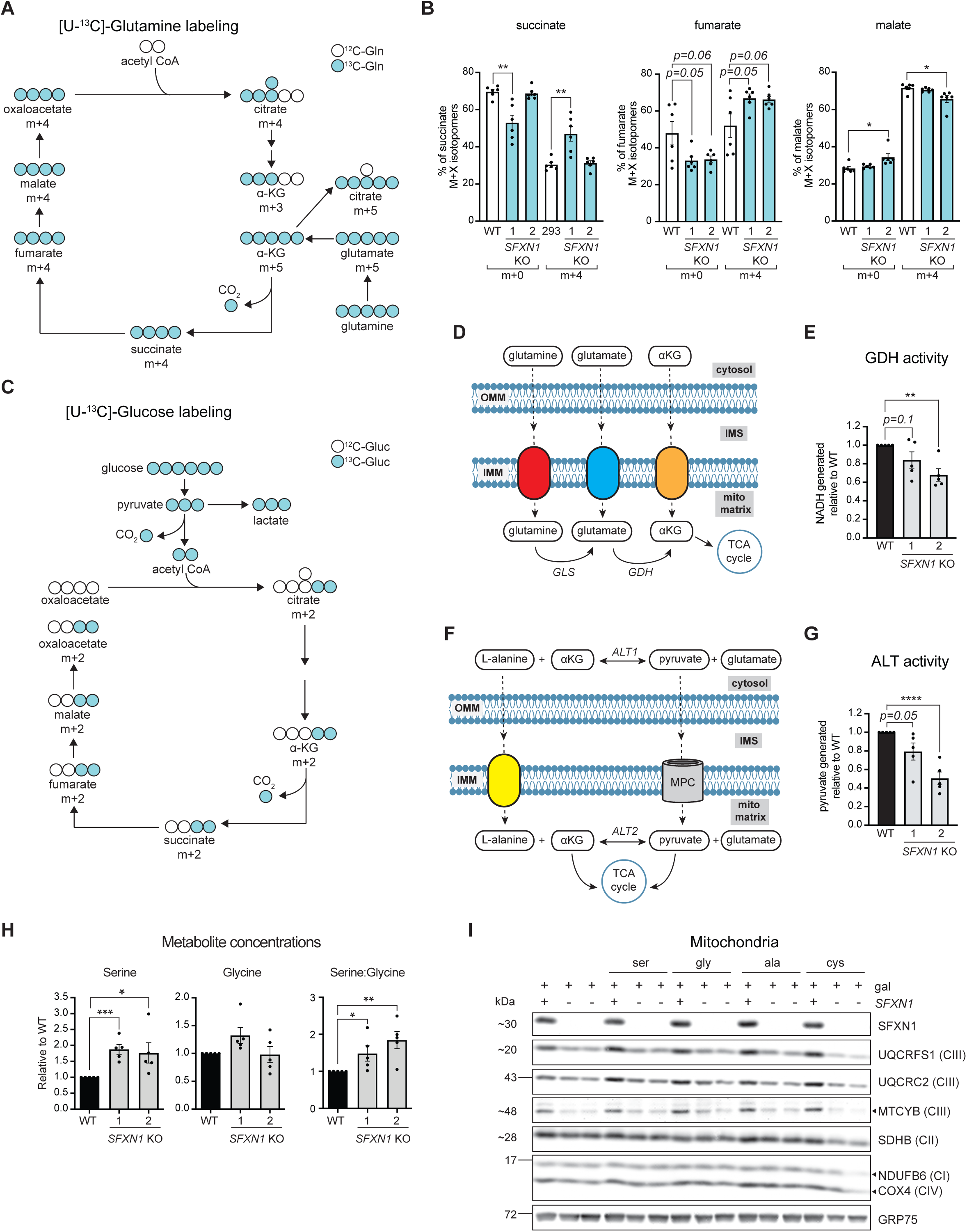
Assessment of central carbon and amino acid metabolism in *SFXN1* KOs. (A) Schematic diagram of [U-^13^C]-glutamine labeling of TCA intermediates. (B) Percent abundance of m+x-labeled TCA intermediates as determined by LC-MS/MS after [U-^13^C]-glutamine labeling (mean±SEM, *n=6*). (C) Schematic diagram of [U-^13^C]-glucose labeling of TCA intermediates (from one turn of the cycle). (D) Reaction mediated by glutamate dehydrogenase (GDH). (E) GDH activity in cellular extracts of HEK293 WT and *SFXN1* KOs. (F) Reaction mediated by alanine aminotransferase (ALT). (G) ALT activity in cellular extracts of HEK293 WT and *SFXN1* KOs. (H) Serine and glycine abundance in cellular extracts relative to WT as determined by fluorimetric-based assays. Serine:glycine ratio is also presented (mean±SEM, *n=5*). (I) Immunoblotting for select respiratory complex subunits using mitochondrial isolates from cells grown for 2 days in galactose-based media only or supplemented with 2 mM serine, glycine, alanine or cystine. \**p<0.05*, \*\**p<0.01*, \*\*\**p<0.001*, unpaired Student’s t-test

**Figure S6 related to Figure 7.**
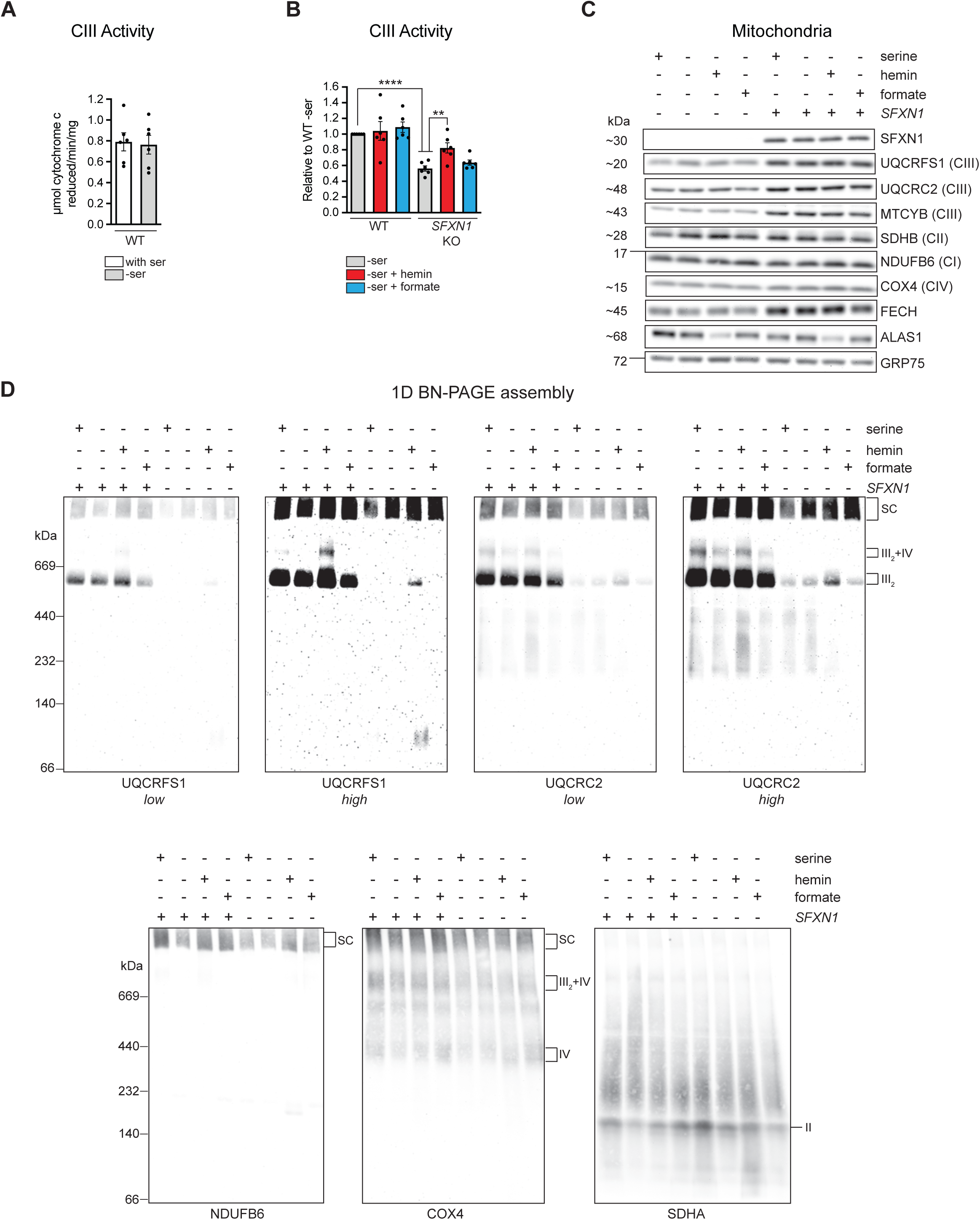
Complex III function in the absence of serine and upon hemin and formate supplementation. (A) Complex III activity in DDM-solubilized mitochondria from cells grown in glucose-containing media with or without serine (mean±SEM, *n*=6). Mitochondrial isolates from cells grown in glucose-containing media without serine, or with addition of hemin or formate were used to assess: (B) Complex III activity (mean±SEM, *n*=6), (C) Steady-state abundance of select proteins, and (D) 1D BN assembly of respiratory complexes. \**p<0.05*, \*\**p<0.01*, \*\*\**p<0.001*, unpaired Student’s t-test

**Figure S7 related to Figure 7.**
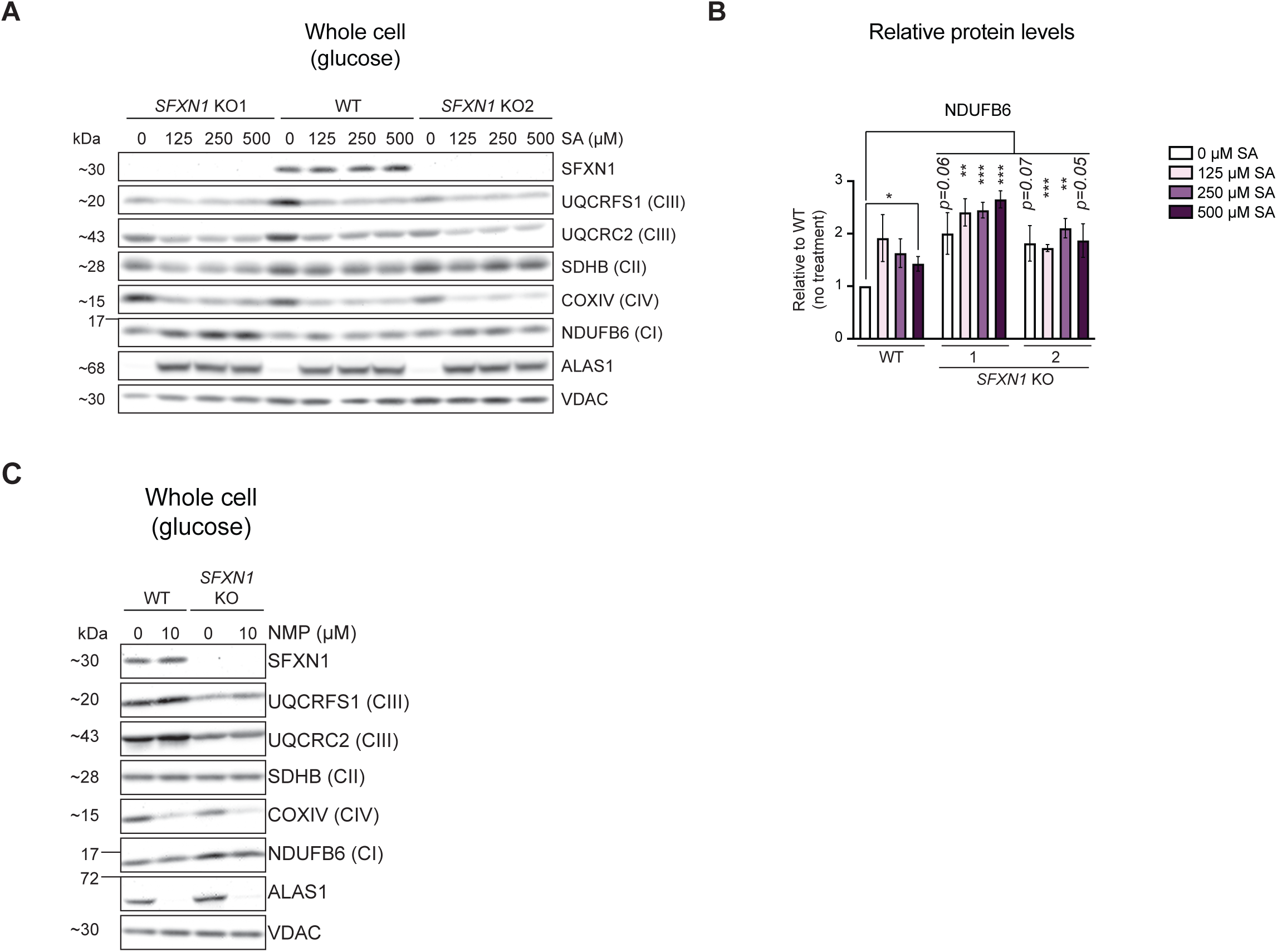
Effect of heme biosynthetic inhibitors on steady-state abundance of respiratory complex subunits. (A) Immunoblotting for select respiratory complex subunits using cell lysates. Cells were grown in glucose-containing media with the indicated SA concentration for 2 days. ALAS1 protein is elevated in response to SA-mediated inhibition of heme biosynthesis. (B) Densitometric analysis of bands in (A) for NDUFB6 and SFXN1. Protein levels in HEK293 WT were set to 1.0. (mean±SEM, *n=3*). (C) Immunoblotting for select respiratory complex subunits using cell lysates. Cells were grown in glucose-containing media with 10 mM NMP for 2 days. ALAS1 protein is decreased in response to NMP-mediated inhibition of heme biosynthesis. \**p<0.05*, \*\**p<0.01*, \*\*\**p<0.001*, unpaired Student’s t-test

